# Regulation of light harvesting in Chlamydomonas: two protein phosphatases are involved in state transitions

**DOI:** 10.1101/2020.03.31.018721

**Authors:** Federica Cariti, Marie Chazaux, Linnka Lefebvre-Legendre, Paolo Longoni, Bart Ghysels, Xenie Johnson, Michel Goldschmidt-Clermont

**Author notes:** Author contributions: MGC and XJ conceived and coordinated the research project, FC, MC, LLL, MGC, PL and BG performed the experiments and analyzed the data, MGC and FC wrote the article with contributions of all the authors, MGC agrees to serve as the author responsible for contact and ensures communication.

## Abstract

Protein phosphorylation plays important roles in short-term regulation of photosynthetic electron transfer. In a mechanism known as state transitions, the kinase STATE TRANSITION 7 (STT7) of *Chlamydomonas reinhardtii* phosphorylates components of light-harvesting antenna complex II (LHCII). This reversible phosphorylation governs the dynamic allocation of a part of LHCII to photosystem I or photosystem II, depending on light conditions and metabolic demands. Little is however known in the green alga on the counteracting phosphatase(s). In Arabidopsis, the homologous kinase STN7 is specifically antagonized by PROTEIN PHOSPHATASE 1/THYLAKOID-ASSOCIATED PHOSPHATASE 38 (PPH1/TAP38). Furthermore, the paralogous kinase STN8 and the countering phosphatase PHOTOSYSTEM II PHOSPHATASE (PBCP), which count subunits of PSII amongst their major targets, influence thylakoid architecture and high-light tolerance. Here we analyze state transitions in *C. reinhardtii* mutants of the two homologous phosphatases, CrPPH1 and CrPBCP. The transition from state 2 to state 1 is retarded in *pph1*, and surprisingly also in *pbcp*. However both mutants can eventually return to state 1. In contrast, the double mutant *pph1;pbcp* appears strongly locked in state 2. The complex phosphorylation patterns of the LHCII trimers and of the monomeric subunits are affected in the phosphatase mutants. Their analysis indicates that the two phosphatases have different yet overlapping sets of protein targets. The dual control of thylakoid protein de-phosphorylation and the more complex antenna phosphorylation patterns in Chlamydomonas compared to Arabidopsis are discussed in the context of the stronger amplitude of state transitions and the more diverse LHCII isoforms in the alga.

## INTRODUCTION

To fulfill their energy requirements, photoautotrophic plants and algae rely on a photosynthetic electron transfer chain embedded in the thylakoid membrane of the chloroplast. Two photosystems (PSII and PSI) and their associated light-harvesting antennae (LHCII and LHCI) mediate the conversion of light energy into chemical energy. In the linear mode of electron flow (LEF), PSII and PSI work in series to extract electrons from water and to reduce ferredoxin (Fd) and NADPH. The two photosystems are connected through the plastoquinone pool, the cytochrome *b*_*6*_*f* complex and plastocyanin. Electron transfer along the chain is coupled to proton accumulation in the luminal compartment of the thylakoid membranes, and the resulting proton gradient is used to drive ATP synthesis. In the cyclic mode of electron flow (CEF), which involves PSI and the cytochrome *b*_*6*_*f* complex, ATP is produced but there is no net reduction of Fd and NADP^+^. The chemical energy that is stored in ATP, reduced Fd and NADPH fuels cell metabolism, and in particular the synthesis of storage compounds such as carbohydrates. In the dark, in sink tissues or during the night, these compounds can in turn provide energy through glycolysis, respiration or fermentation.

While the photosystems are highly conserved in evolution, the light-harvesting antennae and their organization within the photosynthetic supercomplexes, are more diverse. In plants LHCI is constituted of four monomeric subunits (Lhca1-4) stably associated with PSI, while in *Chlamydomonas reinhardtii* (hereafter “Chlamydomonas”), LHCI is composed of ten subunits (two Lhca1 and one each of Lhca2 to Lhca9) (Takahashi et al., 2004; Drop et al., 2011; Mazor et al., 2015; Ozawa et al., 2018; Kubota-Kawai et al., 2019). In plants LHCII is composed of trimers containing combinations of Lhcb1, Lhcb2 and Lhcb3, and of monomeric subunits Lhcb4 (CP29), Lhcb5 (CP26) and Lhcb6 (CP24). In Chlamydomonas, the LHCII trimers are made of 8 different subunits encoded by 9 genes (LHCBM1-9, with LHCBM2 and LHCBM7 sharing an identical amino-acid sequence) while there are only 2 types of monomeric subunits, LHCB4 and LHCB5 ((Elrad and Grossman, 2004; Merchant et al., 2007); reviewed by (Minagawa and Takahashi, 2004; Crepin and Caffarri, 2018)).

Both external and internal factors induce responses that regulate the activity of the photosynthetic machinery (Eberhard et al., 2008). External conditions of temperature and light vary widely, over timescales that range from months for changes linked to the seasons, to minutes or seconds for those related to the weather or the patchy shade of a canopy. Rapid variations in light are challenging for photosynthetic organisms, which have to optimize light harvesting when photon flux is limiting, but avoid photo-damage when light is in excess. The internal demand for ATP or reducing power can also vary widely and rapidly depending on metabolic activities, so that electron transport has to be adapted accordingly through regulatory responses of the photosynthetic machinery (Yamori and Shikanai, 2016).

Under limiting light, the mechanism of state transitions is an important regulator for the redox balance of the electron transport chain, through the reversible allocation of a part of LHCII to either PSII or PSI (Rochaix, 2014). In state 1 (St 1), this mobile part of the antenna is connected to PSII, while in state 2 (St 2) it is at least in part connected to PSI (Nagy et al., 2014; Unlu et al., 2014; Wlodarczyk et al., 2015; Nawrocki et al., 2016; Iwai et al., 2018). State transitions represent a negative feedback regulatory loop that is important to maintain the redox homeostasis of the plastoquinone (PQ) pool: reduction of the pool favors St 2 leading to an increase in the cross-section of the antenna connected to PSI, whose activity oxidizes the pool. Conversely, oxidation of the PQ pool favors St 1 leading to an increase in the cross-section of PSII, which promotes reduction of the pool. Under normal conditions, the system maintains an intermediate state between St 1 and St 2, which ensures a balanced ratio of reduced and oxidized PQ (Goldschmidt-Clermont and Bassi, 2015). In higher plants such as Arabidopsis, state transitions operate under low light in response to changes in light quality, for example under a canopy where the spectrum is enriched in far-red light, which is more efficiently absorbed by PSI than PSII. Thus, St 1 can be experimentally favored by illumination with far-red light, while St 2 is promoted by blue light. In Chlamydomonas, the difference in spectral properties of the two photosystems is not as pronounced (Tapie et al., 1984). However, a strong transition towards St 2 involving a large part of the LHCII antenna is promoted by anaerobiosis in the dark (Wollman and Delepelaire, 1984; Bulte and Wollman, 1990). For lack of oxygen, production of ATP by mitochondrial respiration is prevented, as well as PQH_2_ oxidation by chloroplast PTOX (PLASTID TERMINAL OXIDASE), while fermentative metabolism produces an excess of reducing equivalents leading to a strong reduction of the PQ pool (Bulte et al., 1990; Houille-Vernes et al., 2011). CEF is tuned by redox conditions independently of state transitions (Takahashi et al., 2013). However, by favoring the activity of PSI, St 2 will enhance CEF and the production of ATP, at the expense of LEF and the production of reducing power which depend on PSII (Finazzi et al., 2002).

The transition towards St 2 is regulated through phosphorylation of LHCII by the protein kinase STN7 in higher plants (Bellafiore et al., 2005), or its orthologue STT7 in Chlamydomonas (Fleischmann et al., 1999; Depege et al., 2003; Lemeille et al., 2010). The activation of this kinase requires an interaction with the stromal side of the cytochrome *b*_6_*f* complex and the docking of PQH_2_ to the Q_o_ site of the complex (Vener et al., 1997; Zito et al., 1999; Dumas et al., 2017). The process is rapidly reversible, because the kinase is counteracted by the protein phosphatase PPH1/TAP38 in higher plants (Pribil et al., 2010; Shapiguzov et al., 2010), which favors the transition towards St 1. In Arabidopsis the PSI-LHCI-LHCII supercomplex, which is characteristic of St 2, contains one LHCII trimer which belongs to a pool that would be loosely associated with PSII in St 1 (L trimers) (Galka et al., 2012). This trimer contains both Lhcb1 and Lhcb2 subunits, but it is the phosphorylation of the Lhcb2 isoform which is crucial for state transitions by favoring docking of the LHCII trimer to PSI (Pietrzykowska et al., 2014; Crepin and Caffarri, 2015; Longoni et al., 2015; Pan et al., 2018). Recently, PSI-LHCI-LHCII complexes with a second LHCII trimer have been observed (Benson et al., 2015; Yadav et al., 2017). In Chlamydomonas, STT7 phosphorylates several subunits of LHCII trimers, and also the monomeric antenna LHCB4 (CP29) (Turkina et al., 2006; Lemeille et al., 2010). The other monomeric antenna, LHCB5 (CP26) can also be phosphorylated in this alga (Bassi and Wollman, 1991). In Chlamydomonas, the PSI-LHCI-LHCII complex involves one or two LHCII trimers, and strikingly also the monomeric antennae LHCB4 (CP29) and LHCB5 (CP26) (Takahashi et al., 2006). The LHCBM5 isoform of LHCII was found to be particularly enriched in this complex, but other LHCBM subunits appear to also participate in its formation and to be phosphorylated (Drop et al., 2014). The LHCBM2/7 subunits stabilize the trimeric LHCII and are also part of the PSI-LHCI-LHCII complex, although they may not be themselves phosphorylated (Ferrante et al., 2012; Drop et al., 2014). It has been reported that supercomplexes of PSI involving the cytochrome b_6_f complex can be isolated from cells in St 2 or in anaerobic conditions (Iwai et al., 2010; Steinbeck et al., 2018). They are proposed to facilitate CEF and thus favor the production of ATP. However the presence and significance of these complexes is still a matter of debate (Buchert et al., 2018).

In vascular plants, the protein kinase STN7 has a paralog called STN8 (Vainonen et al., 2005), which is involved in the phosphorylation of numerous thylakoid proteins including the core subunits of PSII (Reiland et al., 2011). The protein phosphatase PBCP is required for the efficient de-phosphorylation of these subunits (Samol et al., 2012; Puthiyaveetil et al., 2014; Liu et al., 2018). While there is some overlap in their substrate specificities, the antagonistic pairs of kinases and phosphatases appear to have fairly distinct roles. STN7 and PPH1/TAP38 are mainly involved in LHCII phosphorylation and state transitions, while STN8 and PBCP influence the architecture of thylakoid membranes and the repair cycle of photo-inhibited PSII. In monocots such as rice, a further role STN8 and PBCP in the high-light induced phosphorylation of the monomeric LHCII subunit Lhcb4 (CP29) has been proposed (Betterle et al., 2015; Betterle et al., 2017; Liu et al., 2018).

While the state-transition kinase STT7 was first identified in Chlamydomonas (Depege et al. (2003), the functional homologs of the plant PPH1/TAP38 and PBCP phosphatases had not been characterized yet in the alga. Here we present evidence that in Chlamydomonas, homologs of PPH1/TAP38 and PBCP play a role in the regulation of state transitions, with partially redundant functions.

## RESULTS

### CrPPH1 plays a role in state transitions

The closest homologue of the Arabidopsis gene encoding the thylakoid phosphatase PPH1/TAP38 (*At4G27800*) was identified in Chlamydomonas as *Cre04.g218150* in reciprocal BLASTP searches (see Materials and Methods) The Chlamydomonas protein, which we call CrPPH1, shares 36% sequence identity and 55% similarity with its *A. thaliana* homologue. In order to investigate the function of CrPPH1 in Chlamydomonas, we obtained a mutant strain from the CLiP library (LMJ.RY0402.16176, (Li et al., 2016)) with a predicted insertion in intron 7 of the *PPH1* gene (Fig. 1A). The site of insertion was confirmed by PCR amplification and sequencing of the amplified product (Fig. S1A). This mutant line showed an alteration of state transitions as will be detailed below. However, according to the CLiP database, this line also carried a second insertion (hereafter *ins2*) in gene *Cre24.g755597.t1.1*, an insertion that was similarly confirmed by PCR analysis (Fig. S1B). To remove the second mutation and to test whether the state transition phenotype co-segregated with the *pph1* mutation, the mutant line (*pph1;ins2*) was back-crossed twice to the parental strain CC5155 (Fig. S1C). From the first cross, 60 complete tetrads were analyzed for paromomycin resistance and for the presence of *pph1* and *ins2* insertions, both of which conferred paromomycin resistance. In 8 of these tetrads where the two insertions segregated (non-parental ditypes with four paromomycin-resistant colonies *pph1 / pph1 / ins2 / ins2*), we found that altered state transitions correlated with the insertion in the *pph1* gene (Fig. S1C). In a second backcross between a *pph1* single mutant deriving from the first backcross and the parental line CC5155, we analyzed 8 complete tetrads and observed co-segregation of the insertion in the *pph1* gene with paromomycin resistance and the state transition phenotype (Fig. S1D). This analysis shows that the alteration in state transitions is genetically tightly linked to the *pph1* mutation. To facilitate biochemical experiments, we further crossed a *pph1* mutant progeny to the cell-wall deficient strain *cw15*, and a double mutant, *pph1;cw15*, was selected for further analysis. All subsequent experiments were performed comparing the parental *cw15* strain with the double mutant *pph1;cw15*, however hereafter for simplicity these strains will be referred to as wild type (WT) and *pph1* respectively. The chlorophyll content and the maximum quantum efficiency of PSII were similar in *pph1* and in the wild type (Table S1). The *pph1* mutant also showed normal growth in a variety of conditions (Fig. S2).

**Fig. 1.**
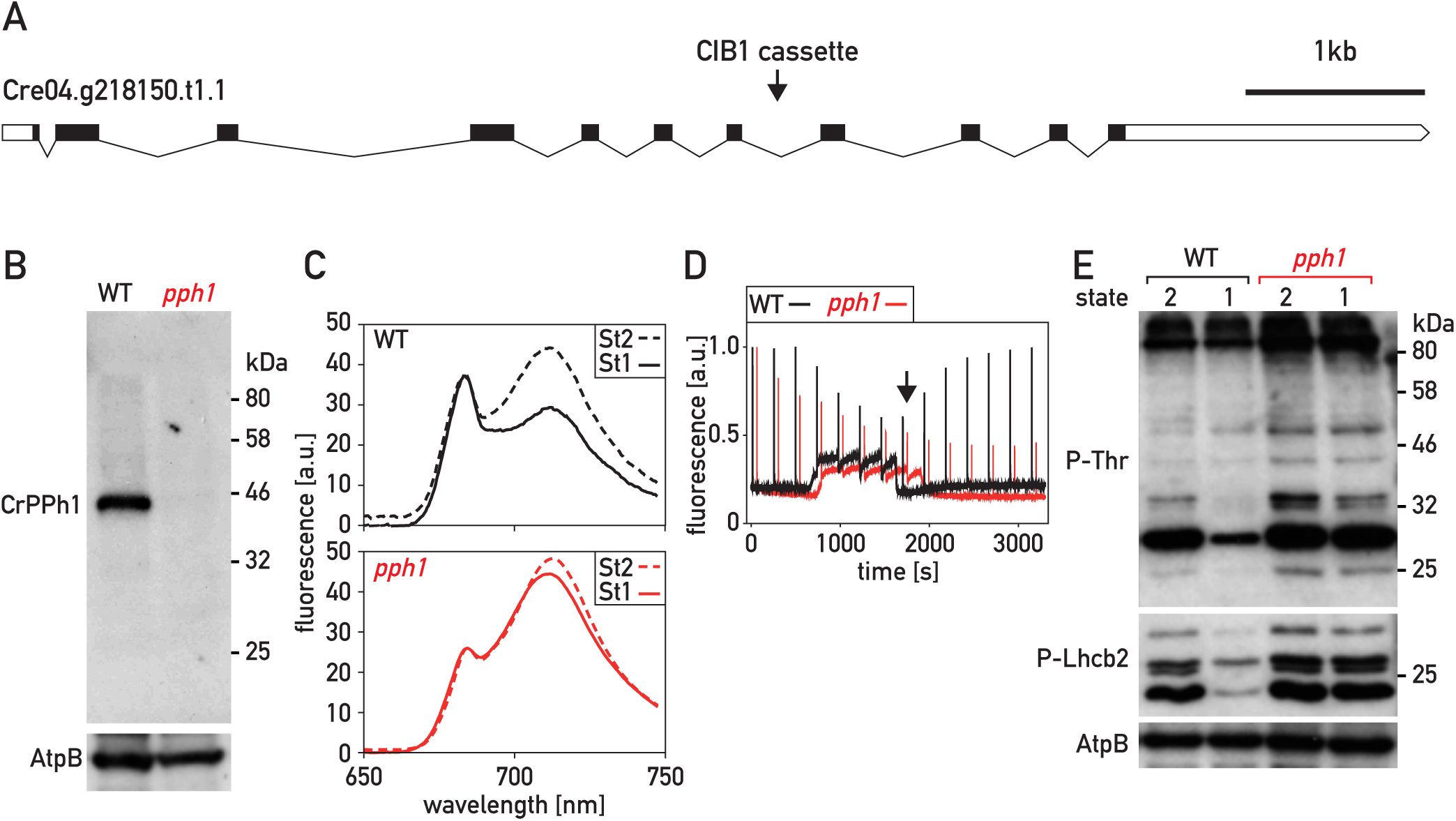
Characterization of the *pph1* mutant. A) Schematic representation of the *PPH1* gene. Exons are represented as black boxes, introns as black lines and the 5’UTR and 3’UTR as white boxes. The arrowhead indicates the site of insertion of the CIB1 cassette in intron 7. B) Immunoblot analysis. Total protein extracts (50 μg) of the wild type (wt) and of the *pph1* mutant were subjected to SDS-PAGE and immunoblotting with antisera against CrPPH1 or AtpB (loading control). C) 77 K chlorophyll fluorescence emission spectra under conditions that favor St 2 (anaerobiosis in the dark) and after 20 min under conditions that favor St 1 (strong aeration). The data are normalized on the PSII peak at 680 nm. D) State transitions monitored by PAM chlorophyll fluorescence spectroscopy at room temperature. Saturating light flashes were fired every four minutes and fluorescence was measured continuously (no actinic light was applied). Data were normalized on the first Fm’ peak. The transition from St 1 to St 2 was induced by sealing the sample chamber to allow respiration to deplete oxygen and cause anoxic conditions, and then the transition from St 2 to St 1 was induced by bubbling air in the sample (at the time indicated with a black arrow). E) Phospho-immunoblot analysis. Total protein extracts (10 μg) of wild type and *pph1* cells in conditions favoring St 2 or St 1 (as in panel B) were subjected to SDS-PAGE and immunoblotting with antisera against P-Thr, P-Lhcb2 or AtpB (loading control).

A rabbit antiserum was raised against recombinant CrPPH1 expressed in *Escherichia coli* (Fig. S3A). It was used to assess the presence of the protein by immunoblotting of total protein extracts of the wild type and of the *pph1* mutant (Fig. 1B). A band was observed in the wild type that was not detected in the *pph1* mutant. This band migrated with an apparent molecular mass of 45 kDa, somewhat slower compared to the calculated 40 kDa molecular mass of CrPPH1 after removal of a predicted chloroplast transit peptide of 52 to 56 amino acid residues according to different algorithms (Predalgo, (Tardif et al., 2012), ChloroP (Emanuelsson et al., 1999)). Because of the background signal with this antibody, a low level of residual CrPPH1 expression in the *pph1* mutant could not be excluded (Fig. S3).

State transitions can be monitored as a change in the fluorescence emission spectrum at 77 K (Fig. 1C). In Chlamydomonas, the transitions can be experimentally induced by switching between anaerobic and aerobic conditions. In the dark, anaerobiosis leads to reduction of the PQ pool and consequent establishment of State 2 (St 2). Subsequent strong aeration under the light promotes oxidation of the PQ pool and a transition to St 1. The relative sizes of the peaks at 682 nm and 712 nm qualitatively reflect changes in the light harvesting antennae associated with PSII and PSI respectively. In the wild type, transition from St 2 (LHCII partly connected to PSI) to St 1 (LHCII mostly allocated to PSII) caused a decrease in the PSI peak relative to the PSII peak. The spectrum was recorded after 20 minutes in the conditions promoting St 1, a timepoint when the transition is nearing completion in the wild type. At this time, the extent of the transition from St 2 towards St 1 was strongly diminished in the *pph1* mutant compared to the wild type (Fig. 1C).

The time courses of state transitions were analyzed in more detail by measuring room temperature Chl fluorescence using Pulse-Amplitude-Modulation (PAM) spectroscopy (Fig. 1D). Cells that had been acclimated to minimal medium in low light (60 μmol photons m^-2^ s^-1^) were transferred to the PAM fluorometer and the sample chamber was sealed, so that consumption of oxygen by respiration led to anaerobiosis. Steady state fluorescence (F_s_) was continuously monitored with the low-intensity measuring beam to monitor the redox state of the PQ pool, and saturating flashes were applied every 4 minutes to measure the maximum fluorescence of PSII (F_m_’), which correlates with the cross-section of its functional light-harvesting antenna. Reduction of the PQ pool in the dark (denoted by an increase in F_s_) triggered the transition to St 2 in the wild type and the *pph1* mutant (decrease of F_m_’). The chamber was then opened and the algal sample bubbled with air, restoring an aerobic environment and allowing the re-oxidation of the PQ pool and a transition towards St 1 (increase of F_m_’). We observed that the transition from St 2 to St 1 was strongly delayed in *pph1* (Fig. 1D), consistent with a role of CrPPH1 in de-phosphorylation of the LHCII antenna. A role in state transitions was confirmed using another protocol (Hodges and Barber, 1983; Wollman and Delepelaire, 1984), whereby the transition from St 2 to St 1 was induced by actinic light in the presence of the PSII inhibitor DCMU (3-(3,4-dichlorophenyl)-1,1-dimethylurea) under continued anaerobiosis (Fig. S4). The fact that at the onset of these experiments, *pph1* is capable of a transition from St 1 towards St 2 implies that although the subsequent transition towards St 1 is delayed, *pph1* can eventually reach at least a partial St 1 under the culture conditions used prior to the measurements (Fig. 1C, D and Fig. S4).

When comparing for the wild type the two protocols (Fig. 1D and Fig. S4A), it is interesting to note that the transition from St 2 to St 1 was rapidly induced by aerobiosis (Fig. 1D), but that its onset was delayed a few minutes when it was induced in the presence of DCMU by actinic light under continued anaerobiosis (Fig. S4A). This difference might be ascribed to a requirement for both an oxidized PQ pool and sufficient levels of ATP to induce the transition towards St 1 (Bulte and Wollman, 1990). While under restored aerobiosis (Fig. 1D), mitochondrial respiration could rapidly replenish the cellular ATP pool, in the presence of DCMU under anaerobiosis (Fig. S4A), cyclic electron flow would replenish the ATP pool more slowly.

To confirm that this state-transition phenotype is due to the *pph1* mutation, we transformed the mutant strain with a plasmid carrying a wild-type copy of *PPH1* in which we inserted a sequence encoding a triple HA (haemagglutinin) epitope at the 3’ end of the coding sequence (*PPH1-HA*) and a selection marker (*aph7”*, hygromycin resistance). The transformants were screened by chlorophyll fluorescence spectroscopy, and 4 lines showing a restoration of state transitions were selected for further analysis. Immunoblotting with a monoclonal HA antibody indicated that the rescued lines (*pph1:PPH1-HA*) expressed the tagged CrPPH1-HA protein, and immunoblotting with the anti-CrPPH1 antibodies showed that the protein was expressed at levels similar to those of the wild type or in moderate excess (Fig. 2A). PAM fluorescence spectroscopy showed that in *pph1:PPH1-HA* the rate of the transition from St 2 to St 1 was comparable to that of the wild type (Fig. 2B).

**Fig. 2.**
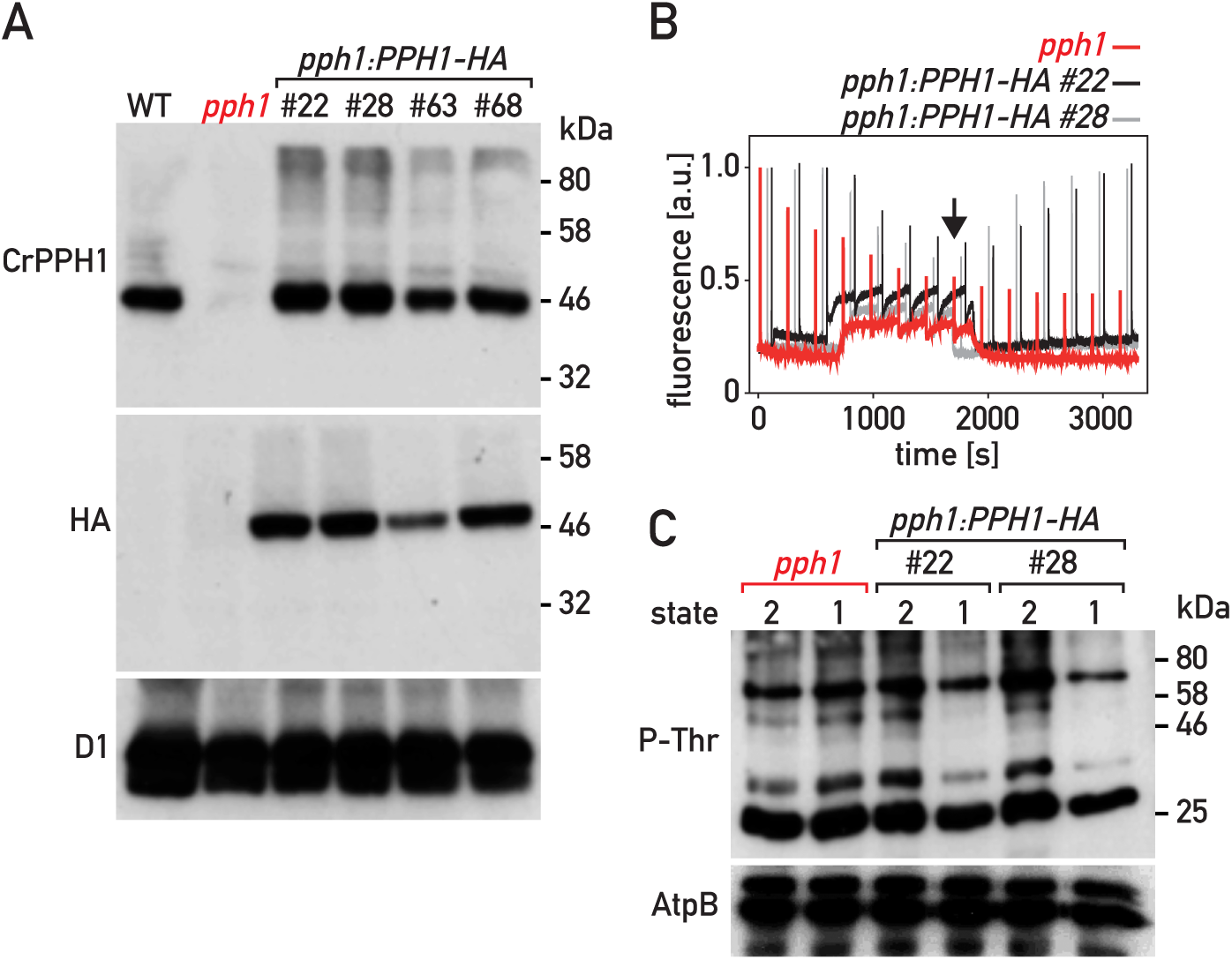
Complementation of the *pph1* mutant. A) Immunoblot analysis. Total protein extracts (50 μg) of the wild type, the *pph1* mutant and four complemented lines (*pph1:PPH1-HA*) were subjected to SDS-PAGE and immunoblotting with antisera against CrPPH1, the HA epitope and D1 (loading control). B) State transitions in the *pph1* mutant and two complemented lines were monitored by PAM chlorophyll fluorescence spectroscopy as in Figure 1D. C) Phospho-immunoblot analysis. Total protein extracts (10 μg) of the *pph1* mutant and of complemented lines were subjected to SDS-PAGE and immunoblotting with antisera against P-Thr or AtpB (loading control).

To further test the role of CrPPH1 in LHCII de-phosphorylation, we examined the phosphorylation status of the major thylakoid proteins by SDS-PAGE and immunoblotting. St 2 was established by anaerobiosis in the dark, and a subsequent transition to St 1 was promoted by strong aeration in the light, as above for the analysis of fluorescence at 77 K. An antibody against phospho-threonine (P-Thr) revealed a complex pattern of phosphoproteins, nevertheless a distinct decrease in the signal of several bands was observed in the wild type upon transition from St 2 to St 1 (Fig. 1E), as well as in the complemented strains (Fig. 2C). As will be shown below, some of the same bands were clearly under-phosphorylated in the mutant *stt7*, which is deficient for the protein kinase involved in state transitions. In the *pph1* mutant, the signal of these phosphoprotein bands decreased much less after switching to the conditions that favor St 1. With an antibody against the phosphorylated form of Arabidopsis Lhcb2 (P-Lhcb2), several bands were observed in Chlamydomonas, corresponding to the migration of LHCII (Fig. S5), although the sequence divergence between the peptide recognized by the antibody and the potential target sequences in the Chlamydomonas antenna subunits does not allow a simple assignment to specific LHCBM isoforms. In the wild type, the intensity of these bands was higher in St 2 than St 1. In the *stt7* mutant (see below) these bands are detected only at low levels that are similar under St 2 or St 1 conditions. These two observations indicate that at least some of the bands revealed in Chlamydomonas by the Arabidopsis anti-P-Lhcb2 antibody are implicated in state transitions. These bands were clearly over-phosphorylated in *pph1* compared to the wild type in the conditions that favor a transition to St 1. Thus, the alteration in state transitions observed spectroscopically in the *pph1* mutant correlates with defects in the de-phosphorylation of LHCII antenna components.

### CrPBCP is also involved in state transitions

A striking feature of the *pph1* mutant was that it showed strong retardation of the transition from St 2 to St 1, but that nevertheless under normal culture conditions it was capable of approaching St 1 and undergoing a subsequent transition to St 2 (Fig. 1D and Fig. S4). It thus appeared that at least one other protein phosphatase might be involved in state transitions and de-phosphorylation of LHCII components, allowing the establishment of St 1 in *pph1*, albeit more slowly. This was corroborated by the identification of another phosphatase mutant affected in state transitions.

A library of mutants with random insertions of an *aphVIII* cassette (paromomycin resistance) was generated and screened using a fluorescence imaging set-up (Tolleter et al., 2011). To search for mutants in photoprotective or alternative electron transfer pathways this library was recently screened again using a different imaging system (Johnson et al., 2009). One of the mutants (identified as *pbcp*, see below) showed impaired transitions from St 2 to St 1 (Fig. 3). Using reverse-PCR techniques, the insertion was mapped to the gene *Cre06.g257850* (Fig. S6A), which encodes the closest homologue in Chlamydomonas of PBCP from Arabidopsis, and will be called CrPBCP hereafter (Fig. 3A). To test whether the alteration of state transitions was genetically linked to the *pbcp* mutation, we backcrossed the mutant to wild type 137C. In 40 complete tetrads, paromomycin resistance segregated with the insertion in the *PBCP* gene. In the 4 tetrads that we analyzed further, an over-phosphorylation of thylakoid proteins co-segregated with the insertion in *PBCP* (Fig. S6B). Moreover, *pbcp* and the phosphorylation phenotype also co-segregated with mating type minus (mt-), as expected because *PBCP* lies in close proximity to the mating-type locus on chromosome 6. One of the *pbcp* mutant progeny was further crossed to *cw15* to obtain a cell-wall deficient strain (*pbcp;cw15*) which was used in all subsequent experiments and will be referred to as *pbcp* hereafter. Using a rabbit antiserum raised against recombinant CrPBCP expressed in *E. coli* (Fig. S3B) and immunoblotting of total proteins separated by SDS PAGE, the protein was detected in the wild type but not in the *pbcp* mutant (Fig. 3B). The chlorophyll content and the maximum quantum yield of PSII were not significantly different in *pbcp* and in the wild type (Table S1). Furthermore, the *pbcp* mutant showed normal growth under a variety of conditions (Fig. S2).

**Fig. 3.**
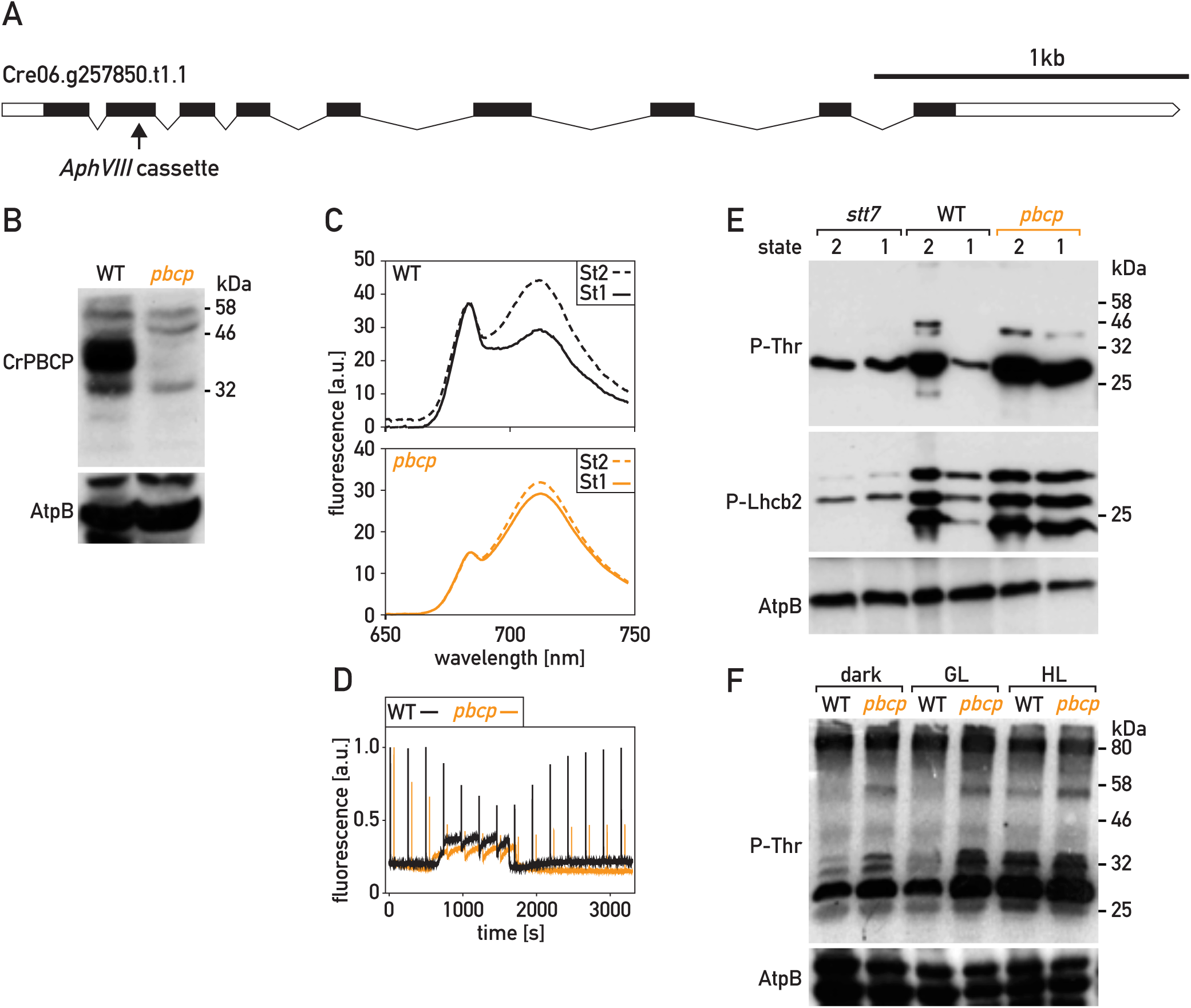
Characterization of the *pbcp* mutant. A) Schematic representation of the *PBCP* gene. Exons are represented as black boxes, introns as black lines and 3’UTR / 5’UTR as white boxes. The arrow indicates the site of insertion of the *aphVIII* cassette in exon 2. B) Immunoblot analysis. Total protein extracts of the wild type and of the *pbcp* mutant (50 μg) were subjected to SDS-PAGE and immunoblotting with antisera against PBCP or AtpB (loading control). C) 77 K chlorophyll fluorescence emission spectra under condition that favor St 2 and after 20 min under conditions that favor St 1, as in Fig 1C. D) State transitions of the wild type (WT) and the *pbcp* mutant were monitored by PAM chlorophyll fluorescence spectroscopy as in Figure 1D. E) Phospho-immunoblot analysis of state transitions. Total protein extracts of wild type and *pbcp* cells in St 2 and St 1 (10 μg, treated as in panel 2 C)) were subjected to SDS-PAGE and immunoblotting with antisera against P-Thr, P-Lhcb2 or AtpB (loading control). F) Phospho-immunoblot analysis of light acclimation. The cells were grown in low light and then transferred to the dark (D), growth light (GL, 80 µE m^-2^ s^-1^) or high light (HL, 300 µE m^-2^ s^-1^) for 2 hours and analyzed like in panel E).

We analyzed state transitions in the *pbcp* mutant by determining its fluorescence emission spectra at 77 K in St 2 and after a subsequent transition to conditions promoting St 1 for 20 minutes. Compared to the wild type, the transition to St 1 was significantly impaired in *pbcp* (Fig. 3C). Using PAM chlorophyll fluorescence spectroscopy, we observed that anaerobiosis in the dark promoted a transition to St 2 in the *pbcp* mutant as in the wild type. However, upon aeration and re-oxidation of the PQ pool, the transition from St 2 to St 1 was strongly delayed in *pbcp* (Fig. 3D). Similar observations were made using the protocol where this transition was induced by light in the presence of DCMU (Fig. S4). Thus, CrPBCP seems to play a major role in state transitions, unlike its homologue in Arabidopsis.

To confirm that the state transition phenotype was due to the *pbcp* mutation, we transformed the *pbcp* mutant with a plasmid carrying a wild-type copy of *PBCP* tagged with a sequence encoding a triple HA epitope (*PBCP-HA*) and a selectable marker (*aph7’’*). Four rescued lines (*pbcp;PBCP-HA*) that expressed the tagged CrPBCP-HA protein were retained for further analysis. Immunoblotting with the anti-PBCP antibodies (Fig. 4A) showed that the different *pbcp;PBCP-HA* lines expressed the protein at levels similar or somewhat reduced compared to the wild type. State transitions were restored in the *pbcp;PBCP-HA* lines, as monitored by PAM fluorescence spectroscopy (Fig. 4B).

**Fig. 4.**
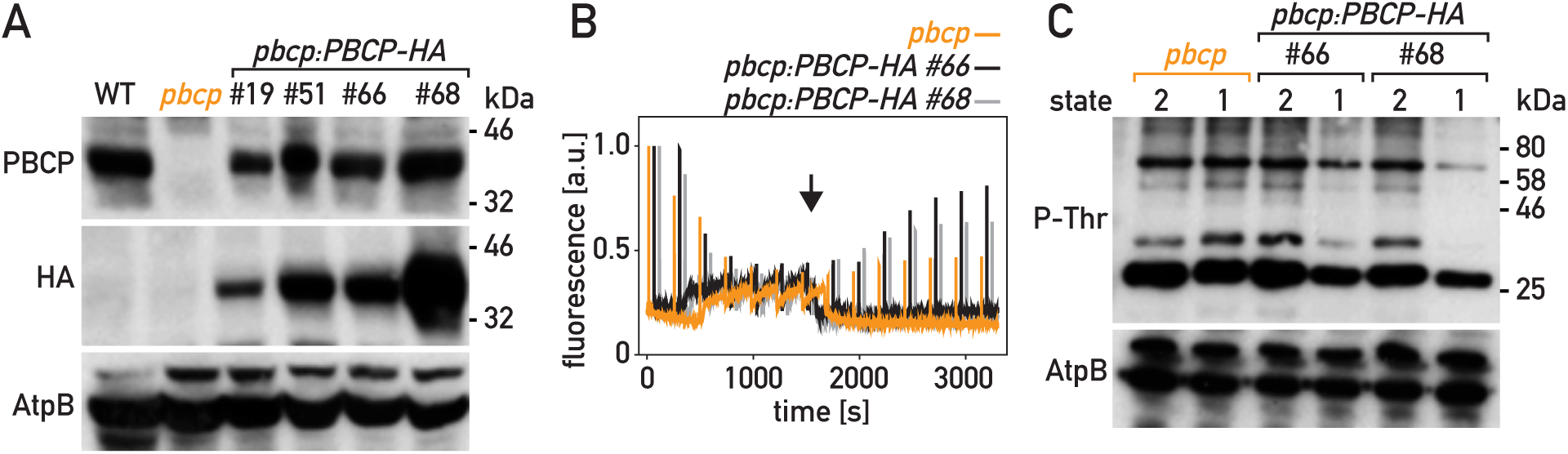
Complementation of the *pbcp* mutant. A) Immunoblot analysis. Total protein extracts (50 μg) of the wild type, the *pbcp* mutant and four complemented lines (*pbcp:PBCP-HA*) were subjected to SDS-PAGE and immunoblotting with antisera against CrPBCP, the HA epitope or AtpB (loading control). B) State transitions in the *pbcp* mutant and two complemented lines were monitored by PAM chlorophyll fluorescence spectroscopy at room temperature as in Figure 1D. C) Phospho-immunoblot analysis. Total protein extracts (10 μg) of the *pbcp* mutant and of complemented lines were subjected to SDS-PAGE and immunoblotting with antisera against P-Lhcb2, P-Thr and AtpB (loading control).

To investigate the alteration of thylakoid protein phosphorylation in the *pbcp* mutant, we used SDS-PAGE and immunoblotting (Fig. 3E). With an anti-P-Thr antibody we observed that bands which migrate as components of LHCII (Fig. S5) were over-phosphorylated in *pbcp*, but not in the complemented *pbcp;PBCP-HA* lines. Unfortunately, these commercial anti-P-Thr antibodies (now discontinued) did not clearly identify the phosphorylated forms of PSII subunits D2 (PsbD) or CP43 (PsbC) (Fig. S5). High phosphorylation of LHCII constituents was confirmed with the Arabidopsis anti-P-Lhcb2 antibodies, which showed over-phosphorylation in *pbcp* of the bands that are under-phosphorylated in the *stt7* mutant. Thus, CrPBCP has a different range of targets in Chlamydomonas than its homologue in Arabidopsis, where the major targets of PBCP are the subunits of the PSII core. It was striking that excess phosphorylation of thylakoid proteins in *pbcp* appeared not only after a transition from St 2 to St 1, but also under normal growth conditions (GL, 80 μmol photons m^-2^ s^-1^), as well as after growth under high light (HL, 300 μmol photons m^-2^ s^-1^) or in the dark (Fig. 3F).

### The *pph1;pbcp* double mutant is locked in state 2

Both CrPPH1 and CrPBCP are implicated in the de-phosphorylation of some components of LHCII and are essential for an efficient transition from St 2 to St 1. Nevertheless, both individual mutants, *pph1* and *pbcp*, are capable of eventually approaching St 1 under the culture conditions prior to the measurements (Figs. 1D, 3D and S4). To further test whether the two phosphatases play partly redundant roles in the regulation of state transitions, we generated double *pph1*;*pbcp* mutants by crossing *pph1* and *pbcp* and genotyping the progeny by PCR (Fig. S7). The *pph1;pbcp* mutants also carry the *cw15* mutation like both their parents. The double mutants were, as expected, deficient for both CrPPH1 and CrPBCP (Fig. 5A). The maximum quantum yield of PSII and the chlorophyll content of *pph1;pbcp* were not significantly different from the wild type (Table S1), and the double mutant showed normal growth under a set of different conditions that were tested (Fig. S2).

**Fig. 5.**
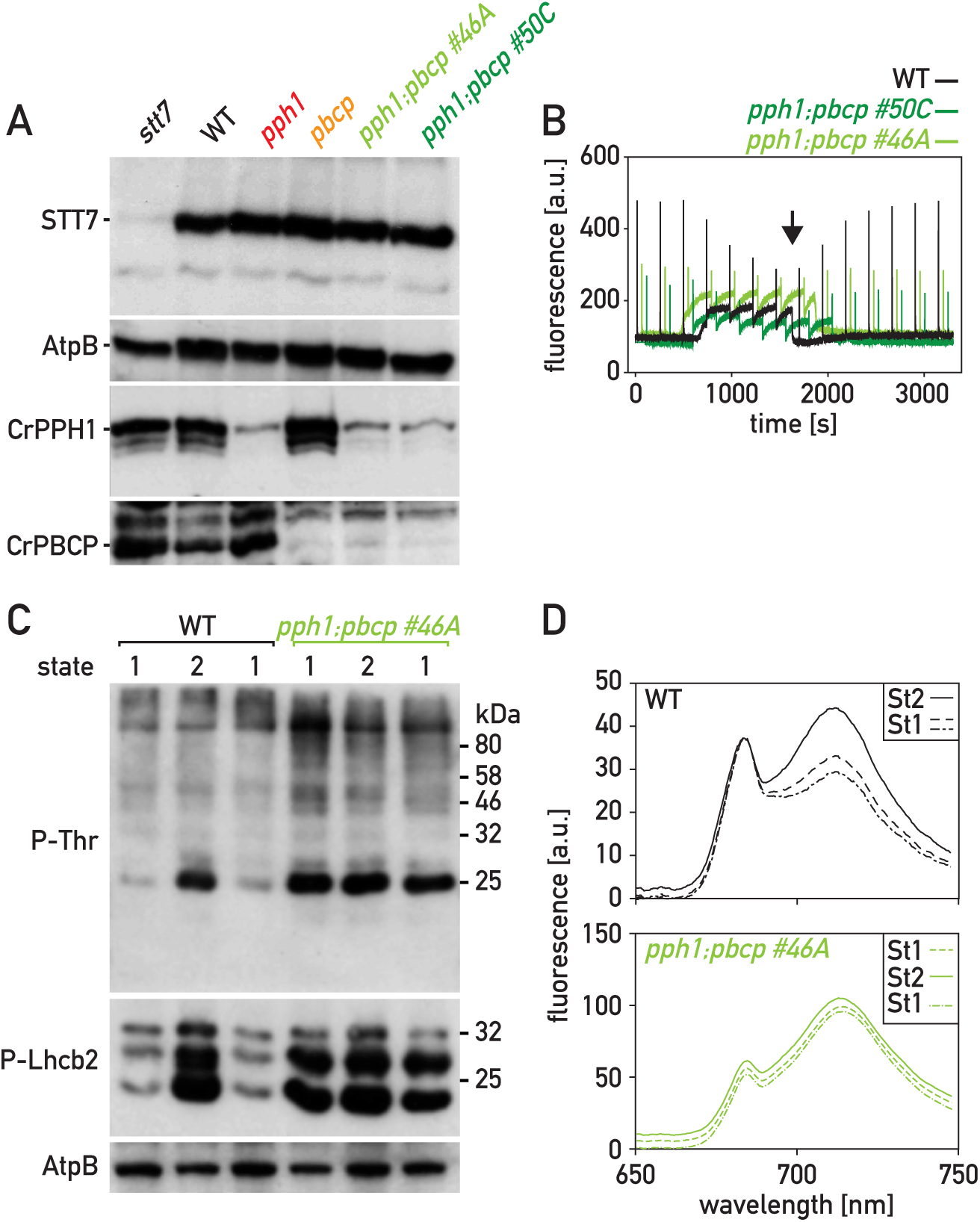
Characterization of the *pph1;pbcp* double mutant. A) Immunoblot analysis. Total protein extracts(50 μg) of the wild type, *stt7, pph1* and *pbcp, pph1;pbcp* (clones # 46A and #50C) were subjected to SDS-PAGE and immunoblotting with antisera against STT7, AtpB (loading control), CrPPH1 and CrPBCP. B) State transitions of the wild type (WT) and two *pph1;pbcp* mutants were monitored by PAM chlorophyll fluorescence spectroscopy at room temperature, as in Figure 1D. The data were not normalized to the first Fm’ peak. C) Phospho-immunoblot analysis. Total protein extracts of wild type and *pph1;pbcp* cells in conditions favoring St 1, then St 2 and finally again St1 (10 μg; cells treated as in panel 1C) were subjected to SDS-PAGE and immunoblotting with antisera against P-Thr, P-Lhcb2 or AtpB (loading control). D) 77 K chlorophyll fluorescence emission spectra under conditions sequentially favoring St 1, St 2 and St 1, obtained as in Fig 1C.

When state transitions were monitored in *pph1*;*pbcp* double mutants, it was remarkable that, following pre-acclimation under low light where the wild-type cells are in St 1, the double mutant had a comparatively low value of F_m_’ at the onset of the measurements, indicative of St 2 (Fig. 5B). There was also no significant further drop in F_m_’ when the *pph1;pbcp* sample became anaerobic with the concomitant rise in F_s_. The double mutant also showed no rise in F_m_’ when the sample was aerated again, indicating that the transition to St 1 was severely hampered. Similar results were obtained when this state transition was promoted by light in the presence of DCMU (Fig. S4). The low level of F_m_’ indicative of St 2 correlated with a high phosphorylation of LHCII components (Fig. 5C). In contrast to the wild type which showed low phosphorylation in St 1, high phosphorylation after a transition to St 2, and again low phosphorylation after a further transition to St 1, *pph1;pbcp* showed in all three conditions a phosphorylation pattern similar to that of the wild type in St 2 (Fig. 5C). These observations indicated that the *pph1;pbcp* double mutant is essentially locked in St 2. In further support of this conclusion, we determined the fluorescence emission spectrum at 77 K of cells in all three conditions (St 1 > St 2 > St 1), and observed that in the *pph1;pbcp* double mutant the fluorescence emission peak of PSI relative to PSII was always larger than in the wild type (Fig. 5 D and Fig. S4B). Our data indicate that two partially redundant protein phosphatases, CrPPH1 and CrPBCP, are involved in the transition from St 2 to St 1.

### Accumulation of thylakoid proteins in the mutants

In Arabidopsis, the expression of STN7 is reduced under prolonged exposure to far-red light (Willig et al., 2011), and the amount of PPH1 is down-regulated at a post-transcriptional level in the *psal* mutant, which is deficient in the docking of LHCII to PSI and thus incapable of completing state transitions (Rantala et al., 2016). These observations prompted us to determine whether in Chlamydomonas the amounts of STT7, CrPPH1 or CrPBCP are altered in the single kinase or phosphatase mutants as well as in the *pph1;pbcp* double mutant, using SDS PAGE and immunoblotting of protein extracts from cells grown under normal conditions (Fig. 5A). However, no significant differences were observed in the accumulation of the three regulatory proteins.

We also investigated the possibility that the phosphatase mutations might be compensated in the long term by changes in the stoichiometry of the photosystems or other major photosynthetic complexes. Proteins extracts of the wild type, the single mutants *pph1* and *pbcp* as well as the double mutant *pph1;pbcp* grown under normal conditions were compared by SDS PAGE and immunoblotting (Fig. S8). Antisera against representative subunits of the major complexes were used for this analysis: AtpB (ATP synthase), D1 (PSII), PsaA (PSI), Cytf (cytochrome b_6_f complex) or COXIIb (mitochondrial cytochrome oxidase). However, no significant differences were apparent in the relative amounts of the photosynthetic complexes in the mutant lines.

### CrPPH1 and CrPBCP have overlapping but distinct de-phosphorylation targets

In Chlamydomonas, LHCII is composed of monomeric LHCB4 (CP29) and LHCB5 (CP26) as well as trimers of isoforms LHCBM1 through LHCBM9. Based on their primary sequences, the trimer subunits belong to four types (Fig. S9): type I in which three subgroups can be distinguished (LHCBM3; LHCBM4/LHCBM6/LHCBM8; LHCBM9), type II (LHCBM5), type III (LHCBM2/LHCBM7, which are identical in mature sequence) and type IV (LHCBM1). While the P-Thr antiserum and the Arabidopsis P-Lhcb2 antiserum showed clear differences in the patterns of LHCII phosphorylation in St 1 in both phosphatase mutants (Fig. 6A), the ill-defined specificity of these antisera did not allow the discrimination of the different components of the antenna, or reveal any target specificity of the respective phosphatases. To address these questions, we used Phos-tag polyacrylamide gel electrophoresis (Phos-tag PAGE) followed by immunoblotting with specific antibodies. We obtained previously described antisera against LHCB4, LHCB5 and LHCBM5 (type II) (Takahashi et al., 2006), and also generated antisera against peptides that are characteristic for the other types of LHCBM subunits. Because these isoforms share a high degree of sequence similarity, the antigenic peptides were selected in the N-terminal region of the proteins, which is the most divergent between LHCII types (Fig. S9). The specificity of the affinity-purified antibodies was tested against the recombinant LHCII subunits expressed in *E. coli* (Fig. S10). The antisera against LHCBM1 (type IV), LHCBM3 (type I), LHCB4 and LHCB5 (minor antenna) proved to be very specific. The antiserum against LHCBM2/7 (type III) showed minor cross-reactions towards type I isoforms. Finally the antiserum against LHCBM4/6/8 (type I, which share the same sequence in the N-terminal region) also reacted towards LHCBM3 (also type I) but unexpectedly very strongly decorated LHCBM9 (type I). It should be noted however, that LHCBM9 is only expressed under conditions of nutrient stress (Grewe et al., 2014).

**Fig. 6.**
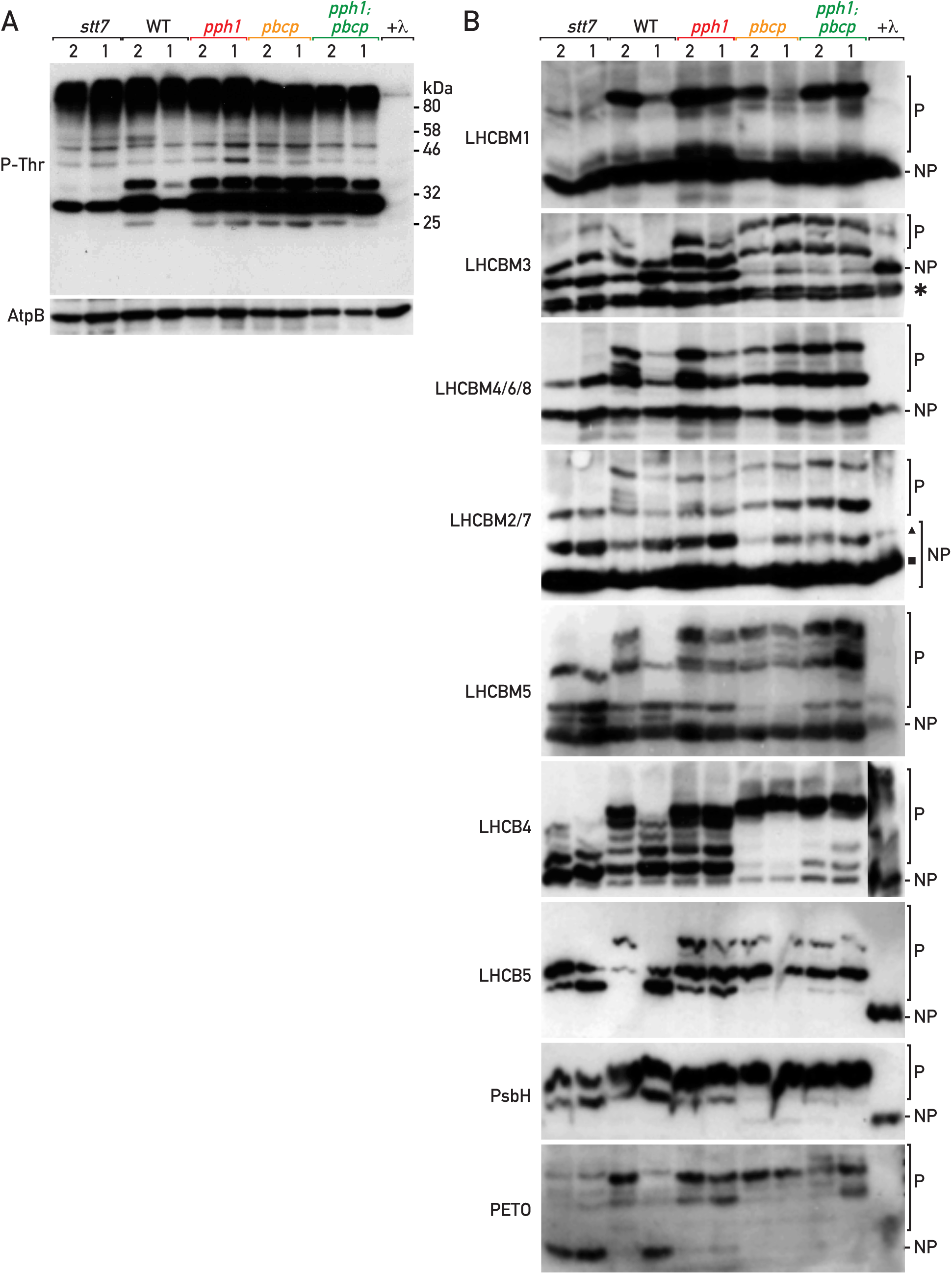
Analysis of CrPPH1 and CrPBCP targets. A) Phospho-immunoblot analysis. Total protein extracts of *stt7*, wild type, *pph1, pbcp, pph1;pbcp* (10 μg) in conditions favoring St 2 and then St 1 were subjected to SDS-PAGE and immunoblotting with antisera against P-Thr or AtpB (loading control). B) Phos-tag PAGE and immunoblot analysis. Total protein extracts (10 μg) of *stt7*, wild type, *pph1, pbcp*, or *pph1;pbcp* in St 2 and then St 1 were subjected to Phos-tag PAGE and immunoblotting with antisera against LHCBM1, LHCBM3, LHCBM4/6/8, LHCBM2/7, LHCBM5, LHCB4, LHCB5, PsbH or PETO. A sample of the wild type in state 2 was treated with lambda phosphatase (+λ) and used as a reference for the migration of the non-phosphorylated form (NP). The migration of the phosphorylated forms (P) is retarded by the Phos-tag immobilized in the polyacrylamide gel. Bands marked with an asterisk, a triangle and a square are discussed in the main text.

To analyze the phosphorylation of the different proteins by immunoblotting, we used Phos-tag which is a metal chelator that can be cross-linked into a polyacrylamide gel during polymerization (Kinoshita et al., 2009). This chelator binds Zn^2+^ ions that interact with phosphate groups and thus retard the migration of phosphorylated forms of proteins during SDS-PAGE. After immunoblotting (Fig. 6B), the degree of phosphorylation of the target protein is reflected in the ratio of the bands representing one or more slower-migrating phospho-form(s) (labelled P) to the band corresponding to the faster-migrating non-phosphorylated protein (labelled NP). To determine the migration of the non-phosphorylated form, we treated a sample of the wild type in St 2 with a non-specific protein phosphatase (λ phosphatase) prior to electrophoresis (lane marked λ in Fig. 6B).The anti-peptide antibodies are targeted to the N-terminus of the LHCBM subunits, which also contains the major sites of phosphorylation. To avoid any bias due to differential recognition by these antibodies of the phosphorylated and non-phosphorylated forms of their targets, we treated the proteins after blotting onto the membrane with λ phosphatase (Longoni et al., 2015). Because the phosphatases play a role in the transition from St 2 to St 1, we analyzed protein extracts of the wild type and the different mutants in St 2 (induced by anaerobiosis in the dark) and after a subsequent transition to St 1 (triggered by aerating the sample).

The antibodies against LHCBM1 (type IV) labelled two bands in the wild type in St 2 (Fig. 6B). The lower one co-migrated with the single band in the λ-phosphatase-treated sample, representing the non-phosphorylated protein. The upper band, corresponding to a phosphorylated form, gave a strong signal relative to the lower one, indicative of a high degree of LHCBM1 phosphorylation in St 2. The ratio of the phosphorylated band to the non-phosphorylated one was much lower in the kinase mutant *stt7*, or after the transition to St 1. In the *pph1* mutant, strong phosphorylation of LHCBM1 was still apparent in the conditions promoting St 1, with a high ratio of the upper to the lower band, in contrast to the wild type or the *pbcp* mutant. The double mutant *pph1;pbcp* showed the same high degree of phosphorylation after the transition to St 1 conditions as the *pph1* mutant. We infer that CrPPH1 is a major actor of LHCBM1 de-phosphorylation during the transition to St 1.

The antibodies against LHCBM3 (type I) decorated two bands after Phos-tag gel electrophoresis of the λ-phosphatase-treated sample (Fig. 6B), as well as after conventional gel electrophoresis of an untreated sample (Fig. S8B), even though these antibodies were specific for LHCBM3 amongst the recombinant proteins expressed in *E. coli* (Fig. S10). The lower band (marked with an asterisk), which was partially resolved as a doublet, is unlikely to represent processed forms of LHCBM3 lacking amino-acid residues at the N-terminus (Stauber et al., 2003) since the LHCBM3 antibodies were raised against this region and would not recognize a truncated protein. Thus the lower band may reflect non-specific binding to another protein. There were two additional slower-migrating bands in the wild type in St 2, largely absent from the λ-phosphatase-treated sample, suggesting that LHCBM3 may undergo phosphorylation at more than one site, or that the non-specific protein is phosphorylated as well. One of these bands were also clearly present in *stt7*, suggesting the involvement of another protein kinase in LHCBM3 phosphorylation. The relative intensity of the top-most of the phosphorylated bands to the non-phosphorylated ones clearly decreased after transition to St 1 in the wild type. Some de-phosphorylation was still apparent in the *pph1* mutant, but the overall phosphorylation level appeared to be higher. In contrast, the *pbcp* mutant showed a much higher ratio of phosphorylated bands in both St 2 and St 1, as did the *pph1;pbcp* double mutant. Thus, CrPBCP appears to play the major role for dephosphorylation of LHCBM3.

With the antibodies against LHCBM4/6/8 (type I), a single major band was observed in the λ-phosphatase-treated sample (Fig. 6B), as well as after conventional gel electrophoresis (Fig. S8B). However, these antibodies also recognized other recombinant type I isoforms, LHCBM3 and LHCBM9 (Fig. S10). The latter should not be expressed under our conditions, since it is induced by nutrient stress (Grewe et al., 2014). After Phos-tag electrophoresis of the wild type sample in St 2, the pattern of phosphorylation was complex, with most bands corresponding to those revealed with anti-LHCBM3 (but excluding the band marked with an asterisk). One of the phosphorylated forms was present in the *stt7* mutant. The degree of phosphorylation of the others decreased in St 1 in both the wild type and *pph1*, but not in *pbcp* or *pph1;pbcp* where the most phosphorylated form was prevalent. We tentatively infer that CrPBCP is involved in de-phosphorylation of LHCBM4/6/8 and, since it also affects LHCBM3, more generally in the de-phosphorylation of type I isoforms.

The antibodies against LHCBM2/7 (type III) decorated a strong band in the λ-phosphatase-treated sample, with a second minor band above it. Likewise, after conventional gel electrophoresis, these antibodies labelled a major and a minor band (Fig. S8B). Amongst the recombinant proteins expressed in *E. coli*, the LHCBM2/7 antibodies strongly recognized LHCBM2/7, but also weakly cross-reacted with the type I isoforms LHCBM3, LHCBM4/6/8 and LHCBM9 (Fig. S10). The patterns of the phosphorylated bands together with the upper non-phosphorylated band (marked with a triangle), but excluding the strong non-phosphorylated band below (marked with a square), were similar to the pattern obtained with anti-LHCBM4/6/8, and may mostly represent the cross-reaction to these isoforms. Compared to LHCBM4/6/8, the strong additional non-phosphorylated band (marked with a square) can tentatively be ascribed to LHCBM2/7, which is thus apparently not subject to phosphorylation under these conditions, or only to a small degree.

With antiserum against LHCBM5 (type II), two bands were detected in the λ-phosphatase-treated sample, and at least five more in the various genotypes in St 2. Two bands were also observed after conventional SDS-PAGE (Fig. S8B), which could represent processed forms of LHCBM5 or cross-reactions to other isoforms, so it is unclear which of the five upper bands correspond to phosphorylated LHCBM5. The four uppermost bands were missing in *stt7* suggesting that their phosphorylation relates to state-transitions. In the wild type in St 1 and in *stt7*, an additional band was present between the two non-phosphorylated ones, likely representing a low-phosphorylation form. Higher ratios of the upper bands to the lower ones were observed in St 1 in *pph1* and *pbcp*, and an even higher ratio in *pph1;pbcp*. These observations suggest that both CrPPH1 and CrPBCP act in a partly redundant manner on the phosphorylation of LHCBM5, and potentially other cross-reacting proteins decorated by the antibody.

Antiserum against LHCB4, which recognized a single band after conventional SDS-PAGE (Fig. S8B), decorated multiple bands in the wild-type in St 2, in accordance with the multiple phosphorylation sites that have been observed in LHCB4 (Lemeille et al., 2010). The upper bands were missing in *stt7*, and the ratio of the upper ones to the lower ones decreased in the wild type in St 1 compared to St 2. In *pph1*, the pattern in St 1 resembled that of St 2, while in *pbcp*, as well as in *pph1;pbcp*, a very high degree of phosphorylation was observed in both states. These data suggest that CrPPH1 and CrPBCP contribute to LHCB4 dephosphorylation in St 1, and that CrPBCP may play such a role also in St 2, independently of state transitions.

The antiserum against LHCB5, which labels a single band after conventional SDS-PAGE (Fig. S8B), decorated at least three bands in the different samples, none of which co-migrated with the non-phosphorylated form in the λ-phosphatase-treated sample. Two of the constitutively phosphorylated forms were present in *stt7*, while the third and slowest-migrating form was present in the wild type in St 2 but not in St 1. The latter was still present in *pph1* in St 1, while it became prevalent compared to the lowest one in *pbcp*, and in *pph1;pbcp*. These observations suggest that CrPPH1 and CrPBCP participate in LHCB5 de-phosphorylation in St 1, and that CrPBCP may also contribute in St 2.

With antiserum against the PsbH subunit of PSII, two phosphorylated forms were observed in the wild type and in *stt7*, in both St 2 and St 1 conditions (Fig. 6B). Phosphorylation appeared stronger in *pbcp* as well as *pph1;pbcp*, suggesting that CrPBCP is probably involved in the de-phosphorylation of PsbH, like its homologue in Arabidopsis (Samol et al., 2012).

Finally with antiserum against PETO, which is a phospho-protein implicated in the regulation of CEF (Hamel et al., 2000; Takahashi et al., 2016), a major phosphorylated form as well as one or two minor ones were observed in the wild type in St 2, that were strongly diminished in St 1 or in the *stt7* mutant. In *pph1* the phosphorylation was retained in St 1, in *pbcp* the upper band became most prevalent, and in *pph1;pbcp*, an even slower-migrating band was apparent. These data indicate that both CrPPH1 and CrPBCP are involved in the dephosphorylation of PETO in a somewhat additive manner.

## DISCUSSION

We have identified two protein phosphatases that are required for efficient transitions from St 2 to St 1 in Chlamydomonas. CrPPH1 is the closest homolog of Arabidopsis PPH1/TAP38, which is involved in state transitions in the plant (Pribil et al., 2010; Shapiguzov et al., 2010). Unexpectedly, we found that in Chlamydomonas CrPBCP is also involved in state transitions. This is in contrast to PBCP from Arabidopsis, which is required for the de-phosphorylation of several PSII core subunits but not of the LHCII antenna (Samol et al., 2012). In Arabidopsis, lack of PBCP does not affect state transitions and it is only when this phosphatase is strongly over-expressed that it has a minor effect on the rate of state transitions. In rice, OsPBCP contributes to de-phosphorylation of LHCB4 (CP29), and is proposed to have a role in regulating the dissipation of excess energy (Betterle et al., 2017). The different substrate specificities of the two phosphatases from higher plants can be explained by the different geometry of their substrate binding sites (Wei et al., 2015; Liu et al., 2018).

The transition from St 2 to St 1 is strongly delayed in the *pph1* and *pbcp* mutants of Chlamydomonas. However after incubation in conditions favoring St 1, both single mutants are nevertheless capable of approaching St 1 (Figs. 1D, 3D and S4), and then to undergo a transition towards St 2. In contrast, the double mutant *pph1;pbcp* remains in St 2 even in the conditions that favor St 1 in the wild type (Figs. 5 and S4). Thus the phenotypes of the two mutants show some additivity, but the two phosphatases have partly redundant functions in state transitions.

State transitions are regulated by the phosphorylation or de-phosphorylation of LHCII antenna proteins. In Arabidopsis, the Lhcb1 and Lhcb2 components of the LHCII trimers can be phosphorylated, but not Lhcb3, with phospho-Lhcb2 playing a major role in the formation of the PSI-LHCI-LHCII complexes in St 2 (Crepin and Caffarri, 2015; Longoni et al., 2015). To investigate the requirement of CrPPH1 and CrPBCP in antenna de-phosphorylation during a transition from St 2 to St 1 in Chlamydomonas, we used Phos-tag PAGE and immunoblotting (Fig. 6B). Although some of the anti-peptide antisera that we developed displayed some cross-reactions amongst closely related LHCBM isoforms, it was still possible to assign many phosphorylated forms to the different types of LHCBM. The Phos-tag PAGE technique has the advantage of allowing an assessment of cumulative phosphorylation. Compared to the simple patterns that are obtained with Lhcb1 and Lhcb2 in Arabidopsis (Longoni et al., 2015), which each one having only a single major phosphorylated form, the patterns proved much more complex in Chlamydomonas, raising the possibility that some LHCBM isoforms may undergo multiple phosphorylation. The analysis revealed interesting differences and overlaps between the targets of CrPPH1 and CrPBCP. CrPPH1 was essential for de-phosphorylation of the type IV isoform LHCBM1, while mainly CrPBCP was required for de-phosphorylation of the type I isoforms LHCBM3 and LHCBM4/6/8. Both CrPPH1 and CrPBCP were necessary for de-phosphorylation of the type II isoform LHCBM5 as well as the minor antennae LHCB4 and LHCB5, with CrPBCP playing a dominant role for LHCB4. An over-phosphorylation phenotype was already apparent in St 2 in the *pbcp* mutant for LHCBM3, LHCB4 and LHCB5 and, although to a lower extent, LHCBM3 and LHCBM5 appeared over-phosphorylated in *pph1* under St 2 conditions. After growth under constant condition of low light or high light, over-phosphorylation was also observed in the *pbcp* mutant by immunoblotting with anti-P-Thr antibodies. It thus appears that CrPBCP may be active in de-phosphorylation status of thylakoid proteins not only in conditions that promote St 1, but also under more balanced steady-state conditions.

In Arabidopsis, PPH1 is specific for LHCII, although its strong over-expression has an effect on the de-phosphorylation of the PSII core subunits D1 and D2 (Pribil et al., 2010; Shapiguzov et al., 2010). Conversely, PBCP is mostly required for dephosphorylation of PSII subunits (D1, D2 and CP43), but it can also act on LHCII (Samol et al., 2012; Longoni et al., 2019). In Chlamydomonas the major phosphorylated thylakoid proteins include, in addition to LHCII, some subunits of PSII (D2 (PsbD), CP43 (PsbC) and PsbH) (Delepelaire, 1984; de Vitry et al., 1991) as well as PETO (Hamel et al., 2000). Using Phos-tag gel electrophoresis, we found that both CrPPH1 and CrPBCP influence not only the de-phosphorylation of components of LHCII, but also of the PsbH subunit of PSII. Phosphorylation of LHCII still occured in a mutant lacking PSII (Wollman and Delepelaire, 1984), but phosphorylation of PSII did not occur in a mutant lacking LHCII proteins (Devitry and Wollman, 1988). While the two phosphatases target partially overlapping sets of components of both PSII and LHCII, it seems likely that differences in regulation and target specificities also lie with the protein kinases STT7 and STL1. STT7 appears specific for LHCII (Turkina et al., 2006; Lemeille et al., 2010), while putatively STL1 might phosphorylate PSII, like its paralog STN8 in Arabidopsis (reviewed by (Rochaix et al., 2012)). We also obtained evidence that both phosphatases are involved in the de-phosphorylation of PETO, which is implicated in the control of cyclic electron flow (Takahashi et al., 2016). In summary, CrPBCP and CrPPH1 seem to have distinct but overlapping sets of targets, with CrPBCP having a somewhat wider spectrum. Whether these targets represent direct substrates of the phosphatases, or indirect targets through regulatory cascades cannot be determined from our genetic analysis.

In Chlamydomonas, LHCBM5, CP29 and CP26 were found to be enriched in PSI-LHCI-LHCII complexes in St 2 (Takahashi et al., 2006; Takahashi et al., 2014), although all types of LHCBM could also be found (Drop et al., 2014). Genetic studies have shown that type III isoform LHCBM2/7, type I isoforms LHCBM4/6/8, as well as CP29 and CP26 are implicated in state transitions (Tokutsu et al., 2009; Ferrante et al., 2012; Girolomoni et al., 2017; Cazzaniga et al., 2020). We observed that, with the possible exception of LHCBM2/7, all types of LHCBMs as well as LHCB4 and LHCB5 are differentially phosphorylated in St 2 compared to St 1. In particular we found that LHCBM1 was phosphorylated in St 2, that its phosphorylation depends on STT7, and that it was de-phosphorylated in St 1, mainly reliant on CrPPH1. However mutants which lack LHCBM1 were previously found to be able to perform state transitions (Ferrante et al., 2012). Taken together these observations beg the question of the physiological role of LHCBM1 phosphorylation. A possible explanation might be that for state transitions LHCBM1 phosphorylation acts redundantly with the phosphorylation of other antenna proteins, so that they can fulfill this role in the mutants lacking LHCBM1. Analysis of knock-down mutants indicated that LHCBM2/7 is important for state transitions (Ferrante et al., 2012). The comparison of the data we obtained with the antibodies against LHCBM2/7 and LHCBM4/6/8, suggests that the type III isoform LHCBM2/7 may not be phosphorylated in St 2, consistently with the previous report that phosphorylation of LHCBM2/7 is not detected in the PSI-LHCI-LHCII complex (Drop et al., 2014). A possible reason might be that LHCBM2/7 could be required structurally for the association of LHCII trimers to PSI-LHCI, but that this isoform would not be involved in regulation through phosphorylation. This would be analogous to the case in Arabidopsis, where Lhcb1 is not phosphorylated when part of the mobile LHCII trimer that binds PSI-LHCI, so that it is only the phosphorylation of Lhcb2 that is required for binding to the PSI docking site (Crepin and Caffarri, 2015; Longoni et al., 2015; Pan et al., 2018).

Compared to Arabidopsis, Chlamydomonas lacks LHCB6 (CP24), but has a more complex complement of the LHCBM subunits composing the LHCII trimers (reviewed by (Crepin and Caffarri, 2018)). Another difference is that in Chlamydomonas, some PSI-LHCI-LHCII complexes contain not only LHCII trimers but also the “minor” antennae LHCB4 and LHCB5 (Drop et al., 2014; Takahashi et al., 2014). Furthermore the amplitude of state transitions is larger in Chlamydomonas than in plants (Delosme et al., 1996), and in the alga strong state transitions are induced by changes in metabolic demands for ATP or reducing power, for example under anoxic conditions (Bulte and Wollman, 1990; Cardol et al., 2009). It remains a matter of speculation whether these differences are related to the more intricate roles of the phosphatases CrPPH1 and CrPBCP in the regulation of light harvesting that have evolved in Chlamydomonas. On the other hand, in Chlamydomonas, phosphorylation of LHCB4 is controlled by CrPBCP, reminiscent of monocots such as rice where phosphorylation of LHCB4 is controlled by OsSTN8 and OsPBCP (Betterle et al., 2017). Further “evo-physio” investigations from a combined evolutionary and physiological perspective on the role of protein phosphorylation in the regulation of photosynthetic acclimation in diverse organisms promise to be a fertile approach.

## MATERIAL AND METHODS

### Strains, growth conditions, and media

The *cw15.16 (mt +)* mutant was obtained by crossing *cw15 (mt -)* to wild type 137C. The *C. reinhardtii* CLiP strain LMJ.RY0402.16176, its parental strain CC4533 (mt -) and the corresponding strain of opposite mating type CC5155 (*mt* +) were obtained from the Chlamydomonas Resource Center (https://www.chlamycollection.org/) (Li et al., 2016). This CLiP strain (*pph1;cao*) was backcrossed twice to CC4533, and then to *cw15* to obtain the cell-wall deficient mutant strain *pph1;cw15* used in this work, and referred to as *pph1*. The *pbcp* strain was isolated by screening random insertional mutants for aberrant chlorophyll-fluorescence induction kinetics. These mutants were generated by inserting an *aphVIII* cassette in wild type 137C (mt-) (Tolleter et al., 2011). The *pbcp* mutant was backcrossed with wild type 137C (mt+), and further to *cw15.16 (mt+)* to obtain the cell-wall deficient mutant strain *pbcp;cw15* used in this work, and referred to as *pbcp*. The genotype of the progeny was verified by PCR (see below) and the phenotype by monitoring state transitions using PAM chlorophyll fluorescence spectroscopy (see below).

Cells were grown in Tris acetate phosphate (TAP) (Harris et al., 1989) under normal growth light (60 - 80 µmol m^-2^ s^-1^) from white fluorescent tubes. The state transitions were obtained with cells in exponential growth phase (2-3 10^6^ cells/mL) collected by centrifugation, resuspended in High Salt Minimal medium (HSM) at a concentration of 2-3 10^7^ cells/mL, and pre-acclimated in dim light (∼10 µmol m^-2^ s^-1^) for 2 h with shaking.

### Mapping of the insertion in *pbcp*

DNA was isolated using the CTAB method and quantified using a NanoDrop spectrophotometer (ThermoScientific). The Resda-PCR protocol was used to identify the left border, the technique was adapted from (Gonzalez-Ballester et al., 2005). The first PCR is performed with Taq polymerase and 10% DMSO. The primers and the program used are described in Table X and Y. A nested PCR was performed on the product of the first PCR diluted to 1/1000, 1/500 or 1/50 using KOD polymerase Xtreme™ (MerckMillipore). The product of the second PCR was then loaded onto an agarose gel 2% and fragments with a molecular weight greater than 800 bp were isolated. PCR products were extracted from the gel and purified with NucleoSpin® Gel and PCR Clean-up Kit (Macherey Nagel) and eluted in water. The Genome Walker technique (Clonetech) was used to identify the right border of the flanking sequence of the cassette insertion site. Genomic DNA was successively digested using the enzyme PvuII followed by the ligation of a specific adapter. The primary PCR uses a primer specific to the insert (AphVIII) and an adapter specific primer. This was followed by a nested PCR whereby products greater than 800 bp are extracted from agarose gel and cloned. Cloning of The PCR products were performed in the vector pGEM®-T (Promega) containing the lactose operon and the gene coding for ampicillin resistance and then transformed into chemocompetent bacteria (DH5) by thermal shock. The bacteria were selected on LB Ampicillin medium in the presence of IPTG (ß-D-1-ThioGalactopyranoside and X-Gal (5-bromo-4-chloro-3-indolyl-beta-D-galactopyranoside) to allow for white/blue selection.

### DNA Constructs

The closest homolog of Arabidopsis thaliana PPH1/TAP38 (At4G27800) was identified in *C. reinhardtii* as Cre04.g218150 in reciprocal BLAST searches performed in Phytozome 12 (E-value: 6. E-57; https://phytozome.jgi.doe.gov/pz/portal.html#!search?show=BLAST) and TAIR 10 (E-value: 9. E-56; http://www.arabidopsis.org/Blast/index.jsp).

Vector pPL18 was obtained from pSL18 by replacing the *aphVIII* resistance cassette with the hygromycin resistance cassette. (Sizova et al., 2001). For this the *aphVIII* cassette was removed by KpnI / XhoI digestion. The hygromycin cassette was amplified from plasmid MAC1_gen3 (Douchi et al., 2016) with primers pPL_56 and pPL_57 (see Table S2) and cloned by Gibson assembly (Gibson et al. (2009)).

The vector was further modified to allow the fusion of a triple HA (hemagglutinin) epitope tag at the C-terminus of the coding sequences of interest. A synthetic fragment encoding the triple epitope (obtained from Biomatik) was cloned into pPL18 by BsmI / NotI digestion, yielding vector pFC18.

The *CrPPH1* coding sequence (CDS) was amplified from a Chlamydomonas cDNA library with primers pFC_216 and pFC_217 and cloned by Gibson assembly into pFC18 linearized with EcoRV / BglII, resulting in pFC18_CrPPH1_HA. The *CrPBCP* CDS was amplified from a cDNA library with primers pFC_218 and pFC_219 and cloned as above into pFC18 resulting in pFC18_CrPBCP_HA. The *CrPPH1* CDS without the predicted chloroplast transit peptide (cTP) was amplified with primer pFC_265 and primer pFC_266 from pFC18_CrPPH1_HA, then cloned into vector pet28a digested with NdeI and SalI to obtain pet28a_CrPPH1_ΔcTP. The *CrPBCP* CDS without the predicted cTP was amplified with primer pFC_267 and primer pFC_268 from pFC18_CrPBCP_HA, and cloned into pet28a as above to obtain pET28a_CrPBCP_ΔcTP. All vectors used were verified by sequencing.

### Transformation

Nuclear transformation by electroporation was modified from (Shimogawara et al., 1998). Cells in exponential phase were collected by centrifugation and resuspended at 10^8^ cell/mL in TAP + 60 mM Sucrose. A volume of 300 µL of cells was incubated with 1 µg plasmid DNA (linearized with XbaI) at 16°C for 20 min, and then the mix was transferred to a 4-mm-gap electroporation cuvette and pulsed at 500 V (C = 50 µF) using a BioRad electroporator (GenePulser II). The cuvette was then incubated at 16°C for 20 min. The cell suspension was diluted into 20 mL TAP, incubated in dim light with gentle agitation for 16h, and collected by centrifugation prior to plating and selection on TAP + 25 μg mL^-1^ hygromycin (Sigma).

### Production of polyclonal antiserum of CrPPH1, CrPBCP, and LHCBMs

For production of CrPPH1 and CrPBCP, recombinant proteins were expressed from plasmids pet28a-CrPPH1_ΔcTP and pET28a-CrPBCP_ΔcTP in *Escherichia coli* BL21(DE3), induced with 1mM IPTG for 4 hours at 37°C. Recombinant proteins purified as in (Ramundo et al., 2013) were used to immunize rabbits and antisera were purified by affinity chromatography with the corresponding protein (Eurogentec).

For production of anti-peptide antisera targeting different LHCBMs isoforms (Natali and Croce, 2015), synthetic peptides (Fig. S9) were used for rabbit immunization and affinity chromatography (Eurogentec). For assessing the specificity and cross-reaction of the antibodies, pET28a plasmids carrying the coding sequences of LHCBM1, LHCBM2, LHCBM3, LHCBM4, LHCBM5, LHCBM6, LHCBM9, CP26 and CP29 (Girolomoni et al., 2017) were used to express all the antenna proteins in *E. coli* BL21(DE3).

### Fluorescence Measurements

Maximum quantum efficiency of PSII (Fv/Fm) of cells that were adapted for ∼1 to 5 min in the dark was measured with a plant efficiency analyzer (Handy PEA; Hansatech Instruments) with the parameters recommended by the manufacturer. State transitions were monitored with a pulse amplitude modulation fluorometer (PAM-Hansatech). A 2 ml sample of cells grown and pre-acclimated in HSM as described above was transferred to the dark in the vessel of a pulse amplitude modulation fluorometer (PAM, Hansatech). The sample was kept in the dark with stirring, and saturating pulses (pulse width of 0.7 sec; intensity 85 %) were applied every 4 minutes. The St 1 to St 2 transition was induced by sealing the chamber, so that respiration led to anoxic conditions, and then the transition from St 2 to St 1 was induced by bubbling air in the sample. The alternate protocol for inducing state transitions in the presence of DCMU is described in Fig. S4. Chlorophyll fluorescence emission spectra at 77 K were measured with a spectrofluorometer (Ocean Optics) with excitation from a LED light source at 435 nm.

### Immunoblotting

For total protein extraction, cells in exponential phase were collected by centrifugation, resuspended in lysis buffer (50 mM Tris-HCl, pH 6.8, 2% SDS, and 10 mM EDTA) and 1× Protease inhibitor cocktail (Sigma-Aldrich) and incubated at 37°C for 30 min with shaking in a thermomixer. Cell debris were removed by centrifugation at 16 000*g* for 10 min at 4°C and the supernatant was used as total protein extract. For protein phosphorylation analysis, any centrifugation step of living cells was avoided by instantly dousing the cultures in 4 volumes of cold acetone. After 30 min incubation on ice and centrifugation at 12 000*g* for 12 min in a SLA-4 rotor (Sorvall), the pellet was treated as before. Samples (10 or 50 μg of total protein) were denaturated for 30 min at 37°C prior to SDS-PAGE (Laemmli, 1970) in 15 or 12% acrylamide gels. After wet transfer the nitrocellulose membranes (Biorad) were blocked in Tris-buffered saline plus Tween (TBST; 20 mM Tris, pH 7.5, 150 mM NaCl, and 0.1% [v/v] Tween 20) supplemented with 5% (w/v) nonfat milk (or 3% (w/v) BSA in case of anti-Phospho-Thr immunoblotting), for 2h at room temperature or 16h at 4°C.

The antisera (and their sources) were as follows: monoclonal anti-HA (Promega), anti-phospho-LHCB2 (Agrisera; AS13-2705), anti-PsaA (a gift of Kevin Redding), anti-Cytf, anti-D1 (gifts of Jean-David Rochaix), anti-Phospho Thr (Invitrogen), anti-LHCB4, anti-LHCB5, anti-LHCBM5 (gifts of Yuichiro Takahashi), anti-rabbit IgG horseradish peroxidase conjugate (Promega), anti-mouse IgG horseradish peroxidase conjugate (Promega). For primary antibody decoration, antibodies were diluted in the same buffers as for blocking and incubation was for 2 hours at room temperature or 16 hours at 4°C. Membranes were washed 4 times for 8 minutes with TBS-T and then incubated 1 hour at room temperature with horseradish-peroxidase conjugated secondary antibody diluted in TBS-T with 5% (w/v) nonfat milk (or 3% (w/v) BSA in case of anti-Phospho Thr immunoblotting). Membranes were washed 4 times for 8 minutes with TBS-T and then revealed by enhanced chemiluminescence (ECL).

### Phos-Tag Gel Electrophoresis

Double layer Phos-tag gels (12 % acrylamide / bisacrylamide 37.5:1; 65 uM Phos Tag) were prepared as in (Longoni et al., 2015) except that the presence of EDTA in the lysis buffer was compensated by adding equimolar Zn(NO_3_)_2_ to the samples, and denaturated for 30 min at 37°C before loading. Protein dephosphorylation on the membranes was performed as in (Longoni et al., 2015). For *in vitro* dephosphorylation, a cell pellet was resuspended in 5mM Hepes pH 7.5, 10 mM EDTA, 1 % TritonX 100; and an aliquot containing 10 µg protein was treated with lambda protein phosphatase reaction mix following the instructions of the manufacturer (New England Biolabs) for 1 h at 30°C.

## ACKNOWLEDGMENTS

This research was funded by the Marie Curie Initial Training Network project, AccliPhot (grant agreement number PITN-GA-2012-316427), the University of Geneva, the Institute of Genetics and Genomics of Geneva (iGE3), the Swiss National Science Foundation (SNF 31003A_146300), the European FP6 program (SOLAR-H Project STRP516510) and the Agence Nationale pour la Recherche (ChloroPaths : ANR-14-CE05-0041-01). M.C. was supported by a scholarship from the Ministry of Science and Education.

We thank Jean-David Rochaix, Jean Alric, Bernard Genty and Gilles Peltier for scientific advice, Pascaline Auroy for technical assistance, Matteo Ballottari for the plasmids expressing LHCBMs, Francis-André Wollman for antiserum against PETO, Yuichiro Takahashi for antisera against CP29, CP26 and LHCBM5, and Nicolas Roggli for help with preparing the figures.

The authors declare no conflict of interest.

## Supplemental Data

**Fig. S1. Genotyping of two insertions in the *pph1* mutant and segregation analysis**

**Fig. S2. Growth properties of the single and double mutants**

**Fig. S3. Validation of CrPPH1 and CrPBCP antisera**

**Fig. S4. Time course of state transitions in the presence of DCMU and corresponding fluorescence emission spectra at 77 K**

**Fig. S5. Migration of selected thylakoid proteins following SDS-PAGE**

**Fig. S6. Identification of the *pbcp* mutant and segregation analysis**

**Fig. S7. Genotyping of *pph1;pbcp* double mutants**

**Fig. S8. Accumulation of photosynthetic proteins in the single and double mutants**

**Fig. S9. Design of peptide antigens for antisera against LHCBM isoforms**

**Fig. S10. Specificity and cross-reactions of the antisera against LHCII components**

**Table S1. Chlorophyll content and maximum quantum yield of PSII in the phosphatase mutants**

**Table S2. List of oligonucleotides used in this work**

