## Supplemental figures and tables for "Regulation of light harvesting in Chlamydomonas: two protein phosphatases are involved in state transitions"

**Fig. S1. Genotyping of two insertions in the *pph1* mutant and segregation analysis**

**Fig. S2. Growth properties of the single and double mutants**

**Fig. S3. Validation of CrPPH1 and CrPBCP antisera**

**Fig. S4. Time course of state transitions in the presence of DCMU and corresponding fluorescence emission spectra at 77 K**

**Fig. S5. Migration of selected thylakoid proteins following SDS-PAGE**

**Fig. S6. Identification of the *pbcp* mutant and segregation analysis**

**Fig. S7. Genotyping of *pph1;pbcp* double mutants**

**Fig. S8. Accumulation of photosynthetic proteins in the single and double mutants**

**Fig. S9. Design of peptide antigens for antisera against LHCBM isoforms**

**Fig. S10. Specificity and cross-reactions of the antisera against LHCII components**

**Table S1. Chlorophyll content and maximum quantum yield of PSII in the phosphatase mutants**

**Table S2. List of oligonucleotides used in this work**

### Figure S1

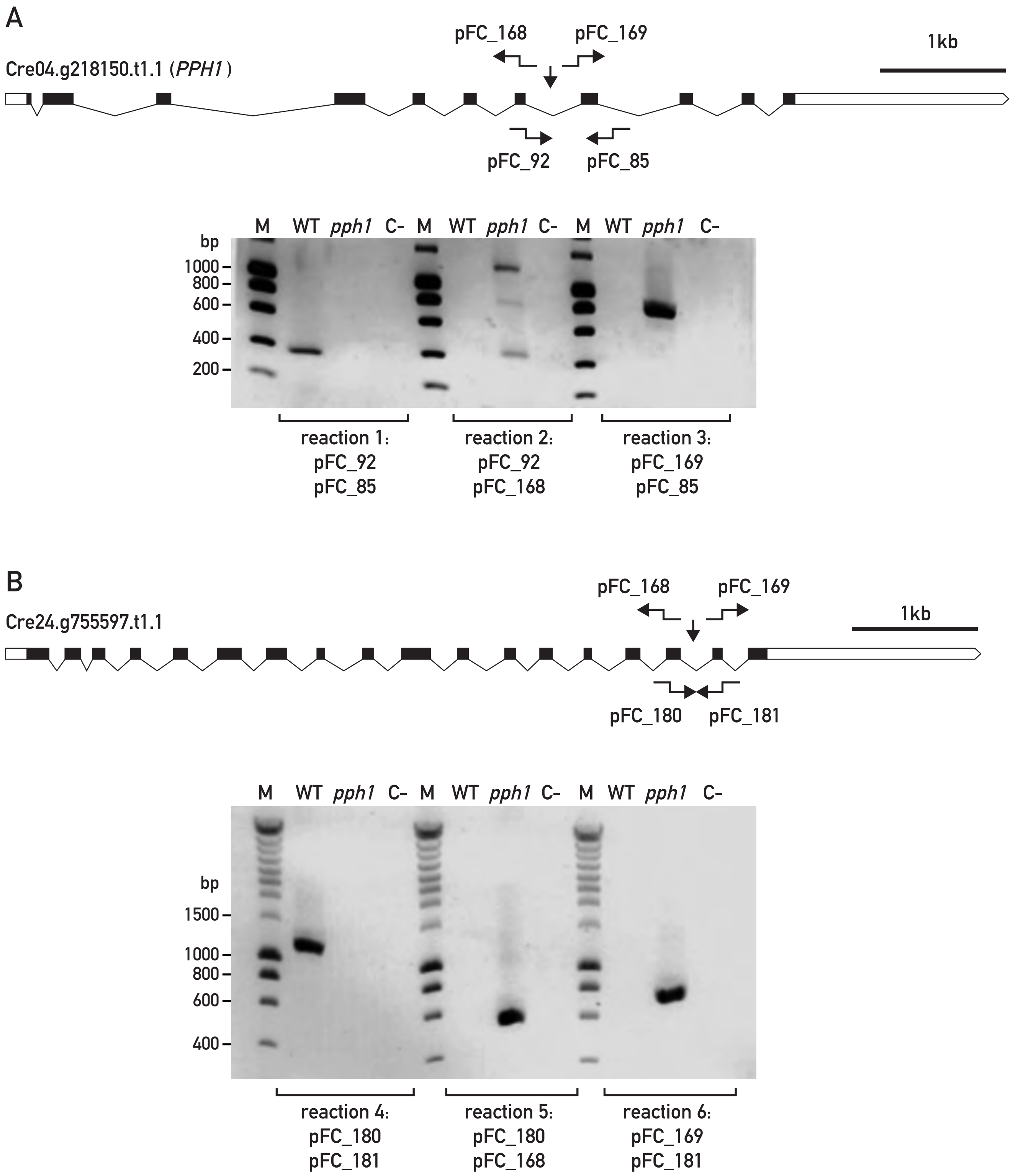


**Fig. S1. Genotyping of two insertions in the *pph1* mutant and segregation analysis**

- PCR mapping of the insertion in the *pph1* gene (*Cre04.g218150.t1.1*). The PCR primers used for genotyping are shown on the gene map and their sequences in table S2.
- Mapping of the second insertion (*ins2*) in gene *Cre24.g755597.t1.1*.
- Segregation analysis of a tetrad from the first backcross. Growth test on TAP + paromomycin, genotyping by PCR of the *pph1* and *ins2* insertions, and state transition phenotypes. Chlorophyll fluorescence to follow state transitions was monitored with a Fluorcam imaging fluorometer (Photon Systems Instruments). Anaerobic conditions that favor St 2 were obtained by flushing with $N_2$ as indicated below the plots (black bars).
- Segregation analysis of a tetrad from the second backcross, as in panel C.

### Figure S1 (continued)

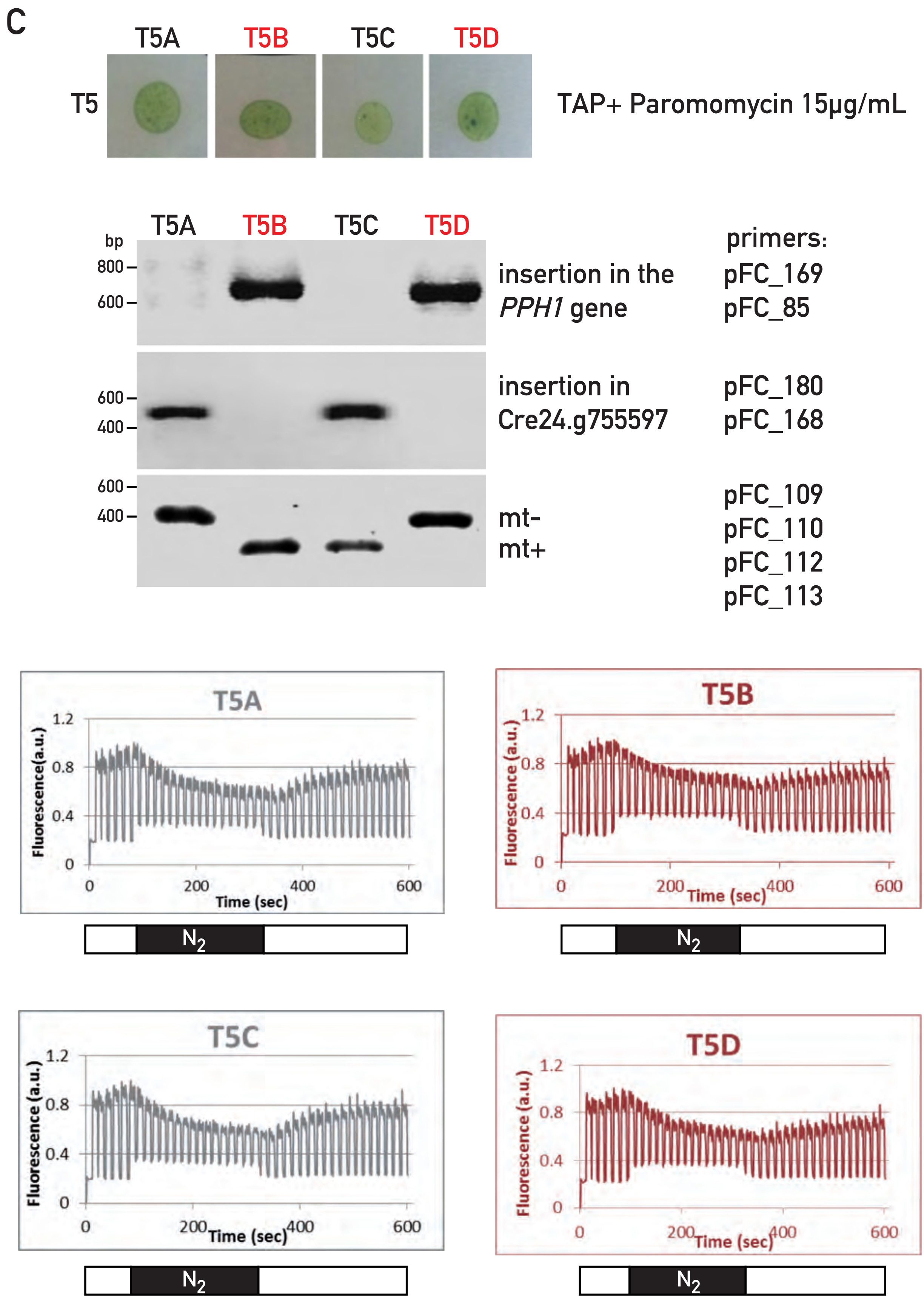

Figure S1 (continued)

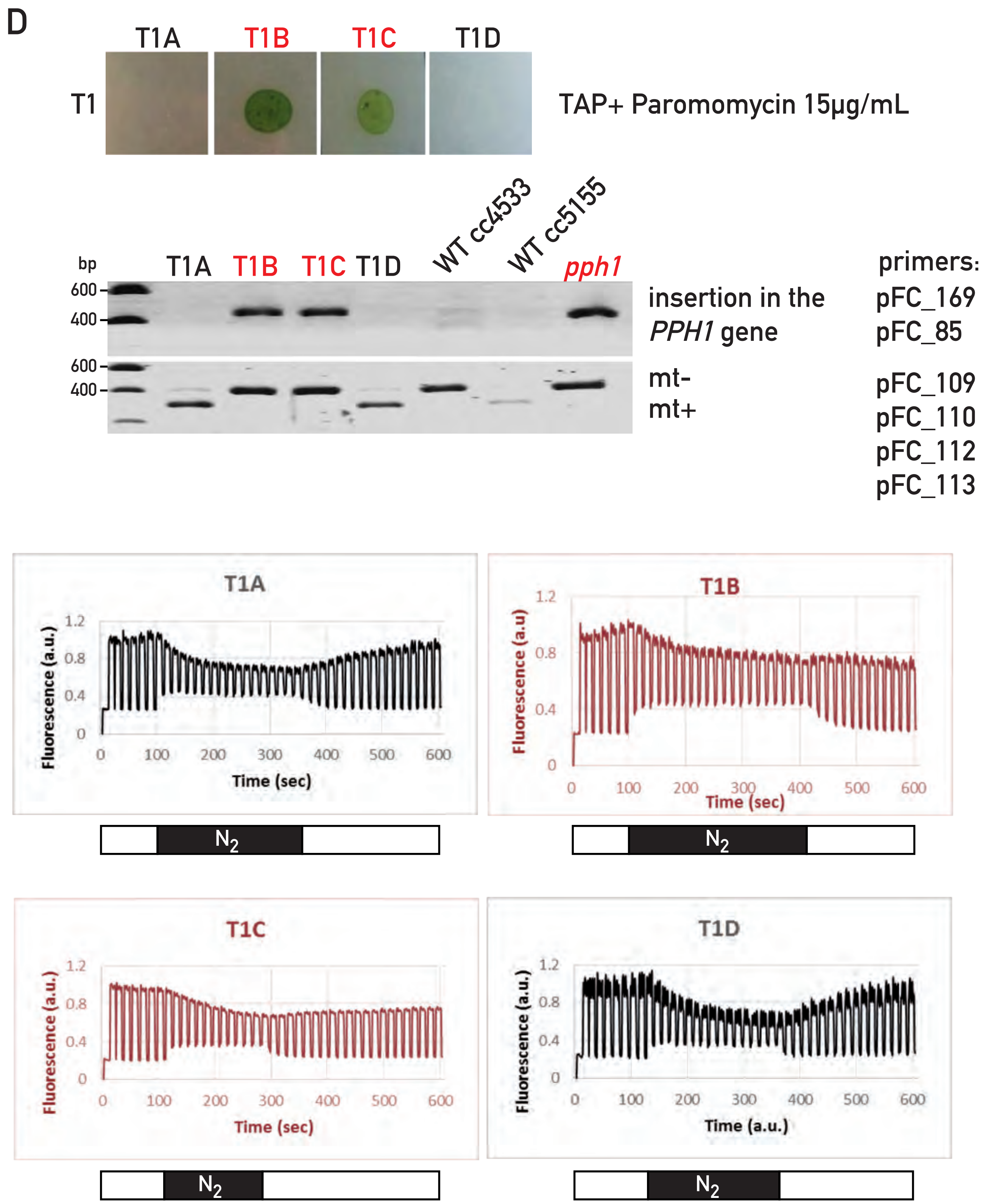

### Figure S2

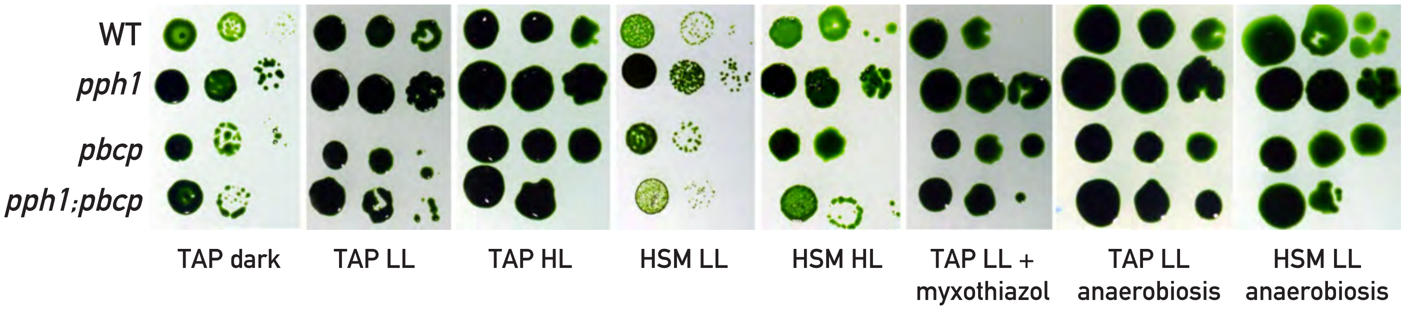


**Fig. S2. Growth properties of the single and double mutants**

Spot tests in different conditions of the wild type (WT) and the *pph1*, *pbcp* and *pph1;pbcp* mutants as indicated on the left. Serial dilutions (10 x) were plated on acetate containing medium (TAP) or on minimal medium (HSM), under low light (LL; 60 µmol photons $m^{-2}$ $s^{1}$) or high light (HL, 250 µmol photons $m^{-2}$ $s^{1}$). Where indicated, anaerobiosis was obtained with Anaerocult P bags (Merck),or myxothiazol was included in the medium (5 µM, an inhibitor of mitochondrial respiration, previously shown to inhibit growth of the *stt7* mutant (Cardol *et al.* 2009)).

### Figure S3

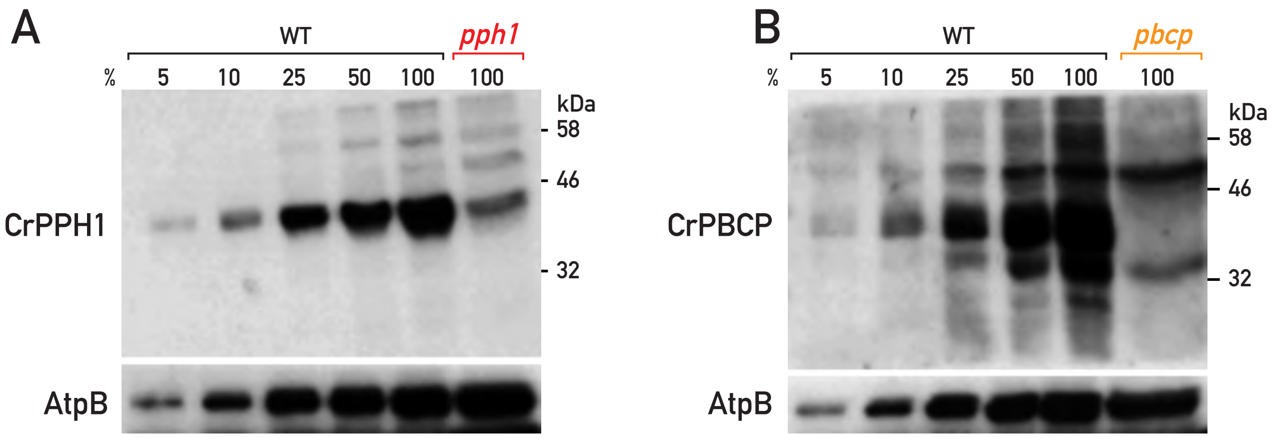


**Fig. S3. Validation of the CrPPH1 and CrPBCP antisera.**

A. Immunoblot with antiserum against CrPPH1 on total protein extracts from *pph1* and the wild type (WT, loaded in different amounts, 100 % corresponds to 100 µg of protein). AtpB is used as loading control.

B. Immunoblot with antiserum against CrPBCP on total protein extracts from *pbcp* and the wild type (WT), as in panel A.

### Figure S4

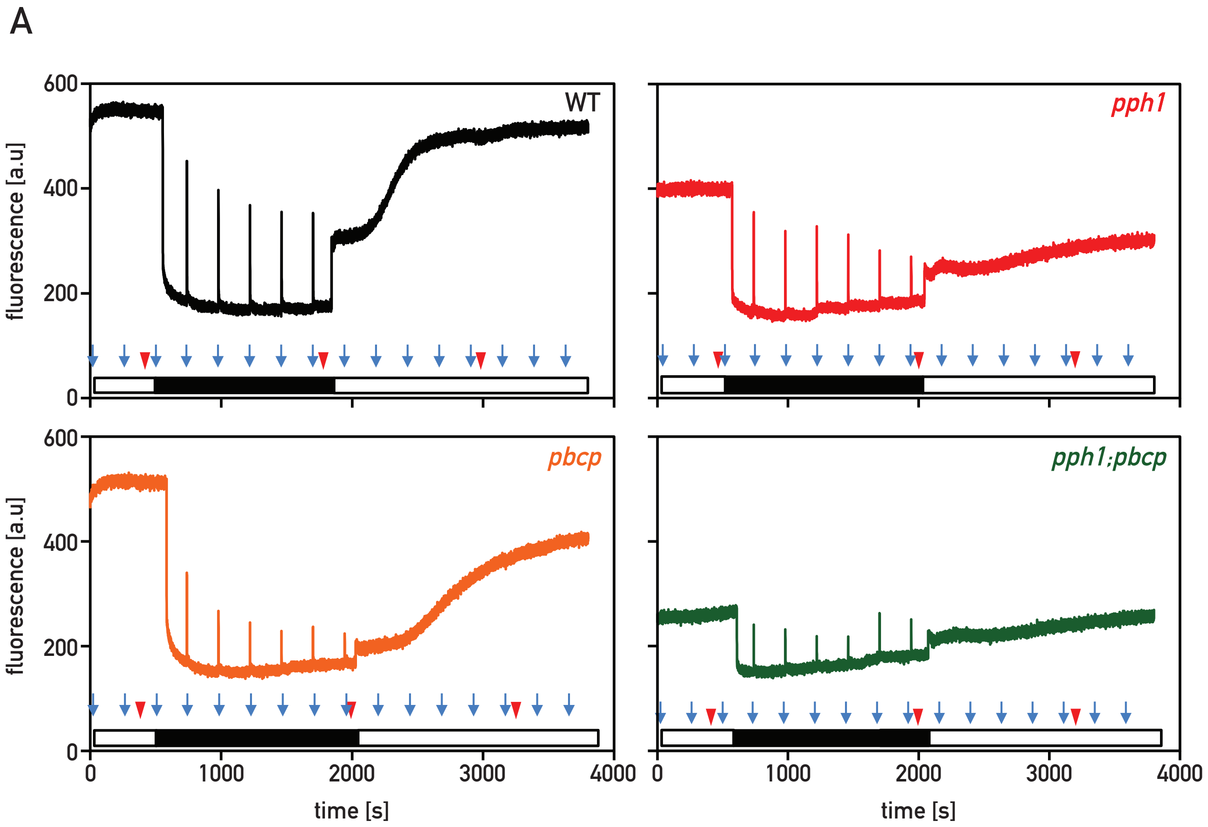


**Fig. S4. Time course of state transitions in the presence of DCMU and corresponding fluorescence emission spectra at 77 K.**

A. State transitions were monitored using chlorophyll fluorescence with a pulse amplitude modulation fluorometer (PAM-Hansatech). Cells grown and pre-acclimated in HSM as described in Material and Methods (2 mL, 2 $10^7$ cells/mL) were transferred to the fluorometer chamber and DCMU (20 µM) was added (Hodges and Barber, 1983; Wollman and Delepelaire, 1984). Saturating pulses (0.7 sec) were applied every 4 minutes (shown with blue arrows). The sample was illuminated with white actinic light (white bar, 400 µmol $m^{-2}$ $s^{-1}$) followed by a dark period (black bar). The chamber was sealed at the beginning of the dark period so that consumption of oxygen by respiration led to anaerobiosis and reduction of the PQ pool. Subsequently the cells were illuminated again with the same intensity white actinic light (white bar).
In the wild type and single mutants, the transition towards St 2 was induced by anaerobiosis as seen by the reduction in PSII yield in the dark. When the cells were returned to light, the activity of PSI with the concurrent block of PSII by DCMU led to the oxidation of the PQ pool and induction of the transition from St 2 to St 1, which was clearly observable in the wild type as a rapid increase in the fluorescence yield of PSII. This increase was delayed in the *pbcp* mutant, even more so in *pph1*, and hardly detectable in the double mutant *pph1;pbcp*.
The curves are representative of 3 (WT, *pph1*, *pbcp*) or 2 (*pph1;pbcp*) independent biological replicates. Red arrowheads indicate the times when samples were taken for the measurement of fluorescence emission spectra (panel B).

B. State transition kinetics were concomitantly monitored by taking samples from the chamber of the PAM fluorometer and determining their chlorophyll fluorescence emission spectra at 77 K (see Material and Methods). The initial spectrum under St 1 conditions (dotted line) was measured in the light, the spectrum under St 2 conditions (dashed line) was measured after 20 - 30 minutes in the dark, and the transition from St2 to St 1 (continuous line) was monitored after exactly 20 minutes of light exposure. At this time point the transition to St 1 is nearly complete in the wild type. The data are normalized on the PSII peak at 680 nm.

### Figure S4 (continued)

B

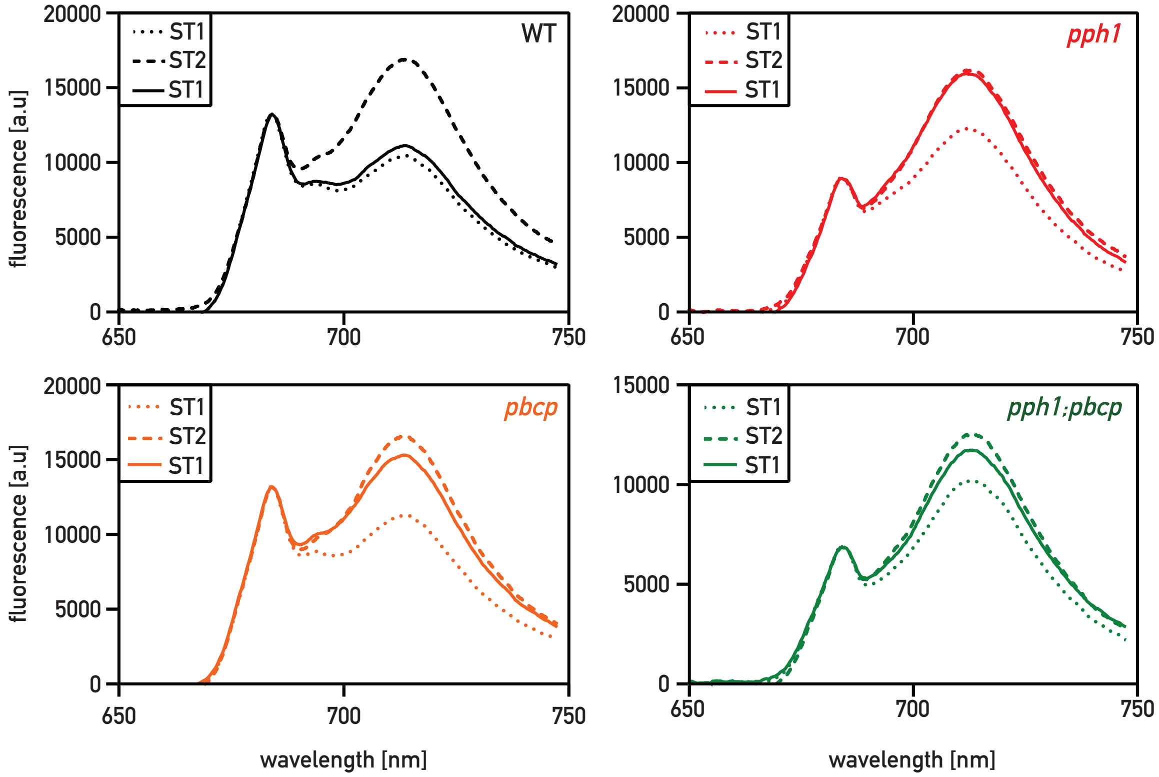

### Figure S5

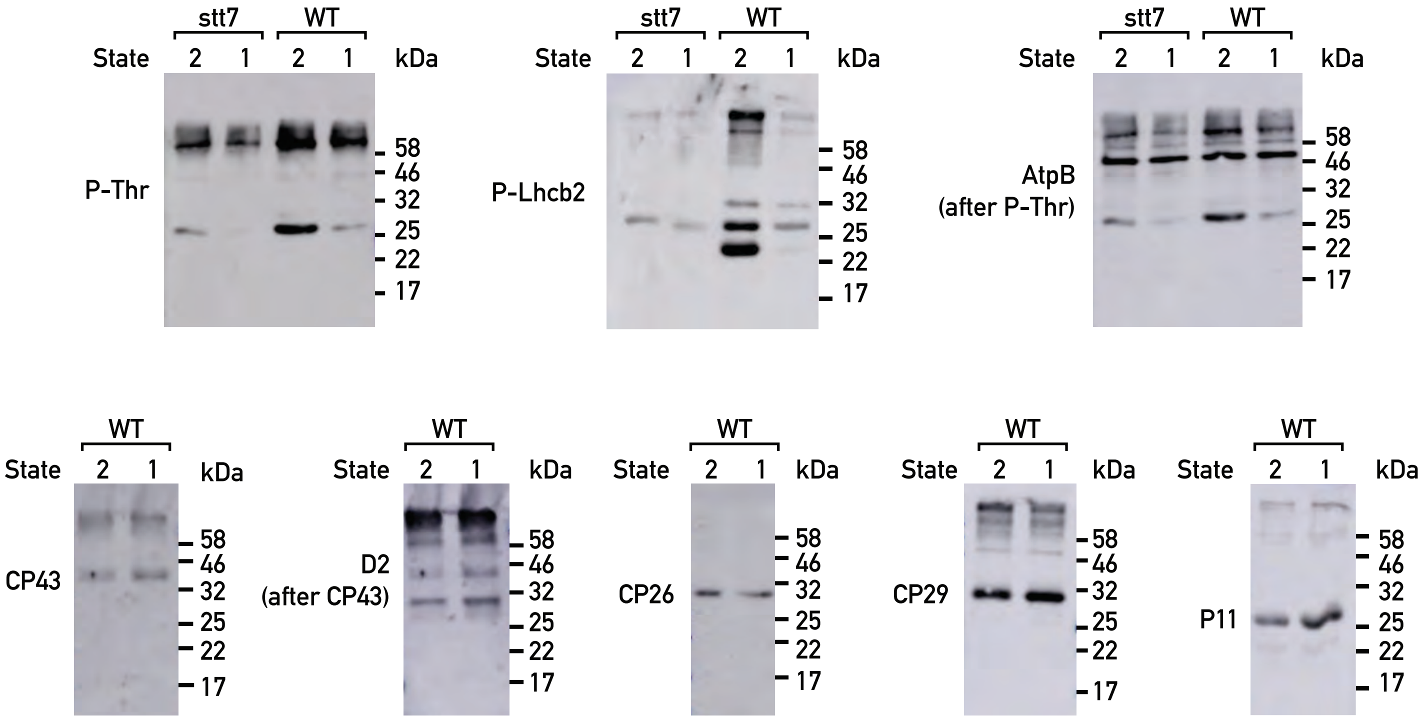


**Fig. S5. Migration of selected thylakoid proteins following SDS-PAGE.**

Total protein extracts (10 µg) of wild type and *stt7* cells in conditions favoring St 2 or St 1 (as in Fig. 1C) were subjected to SDS-PAGE and immunoblotting with antisera against P-Thr, P-Lhcb2, AtpB (reprobed after P-Thr), CP43, D2 (reprobed afer D2), CP26, CP29 and P11 (LHCII).

### Figure S6

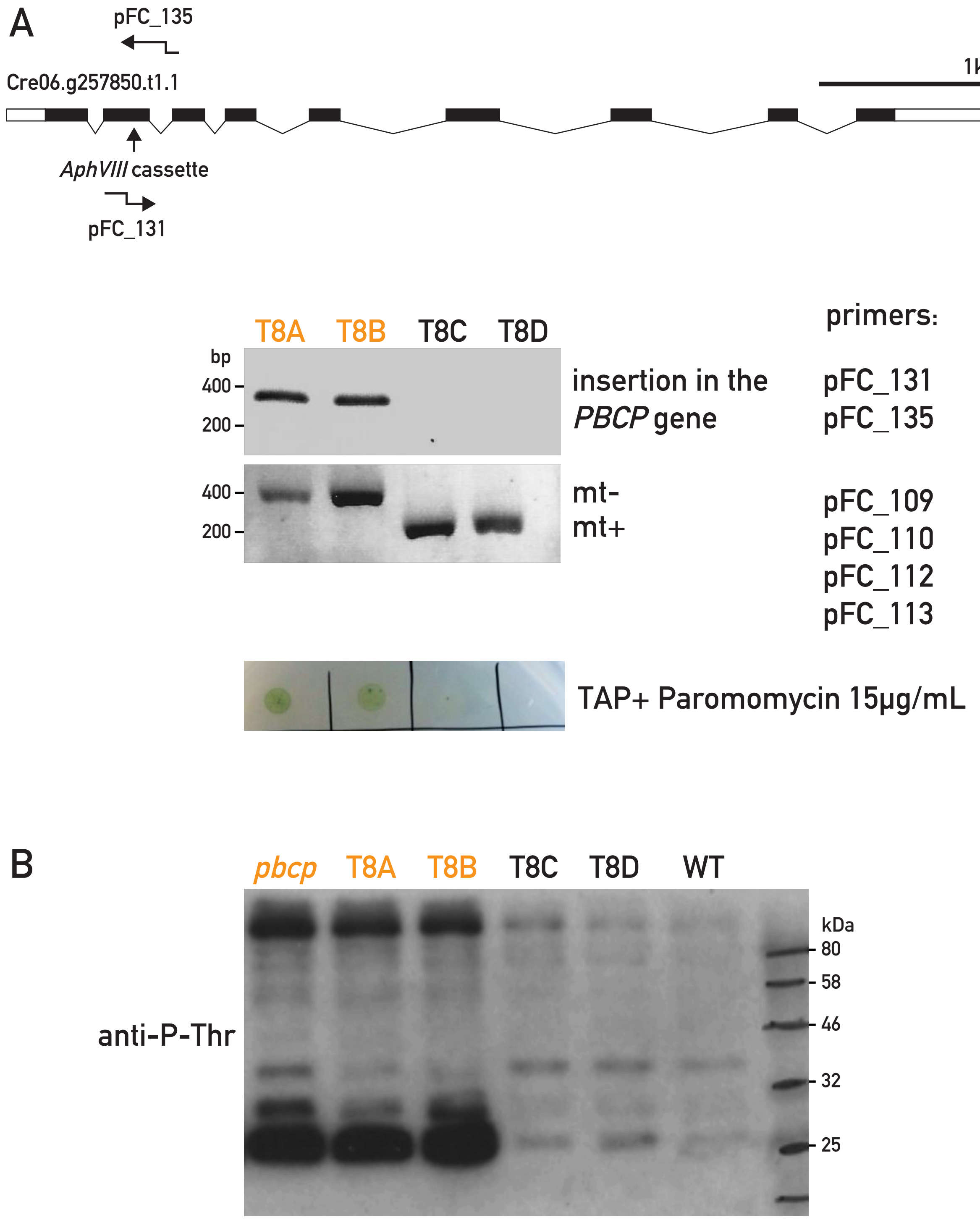


**Fig. S6. Identification of the *pbcp* mutant and segregation analysis**

A. Map of the insertion site in *pbcp* in the gene model *Cre06.g257850*. Two fragments of the pSLX cassette (*AphVIII*) were mapped to exon 2 (466 base pairs from the predicted beginning of the transcript 5'UTR). The left border was determined using the RESDA technique and the right border using the Genome Walker technique. The insertion is not accompanied by deletion or rearrangement of *PBCP* in this region.

B. Segregation analysis. The linkage analysis of a representative tetrad from the first backcross is shown. Genotyping by PCR, growth tests on TAP + Paromomycin and phospho-protein analysis by immunoblotting with P-Thr antiserum (cells were collected from a culture in HSM under low light) are presented.

### Figure S7

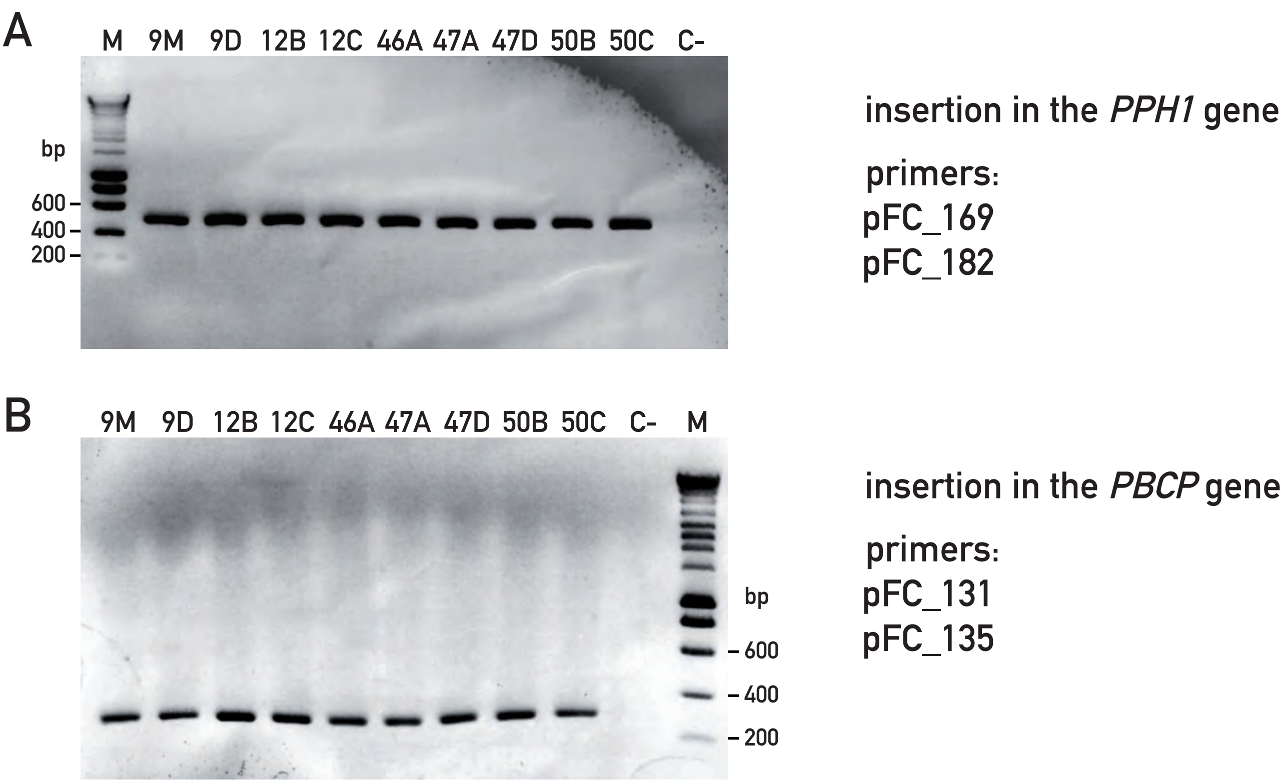


**Fig. S7. Genotyping of *pph1;pbcp* double mutants.**

A. Genotyping by PCR of the insertion in the *PPH1* gene in nine different clones of *pph1;pbcp* double mutants. The PCR primers used for genotyping are shown, and their sequences listed in Table S2.

B. PCR mapping the insertion in the *PBCP* gene as in panel A

### Figure S8

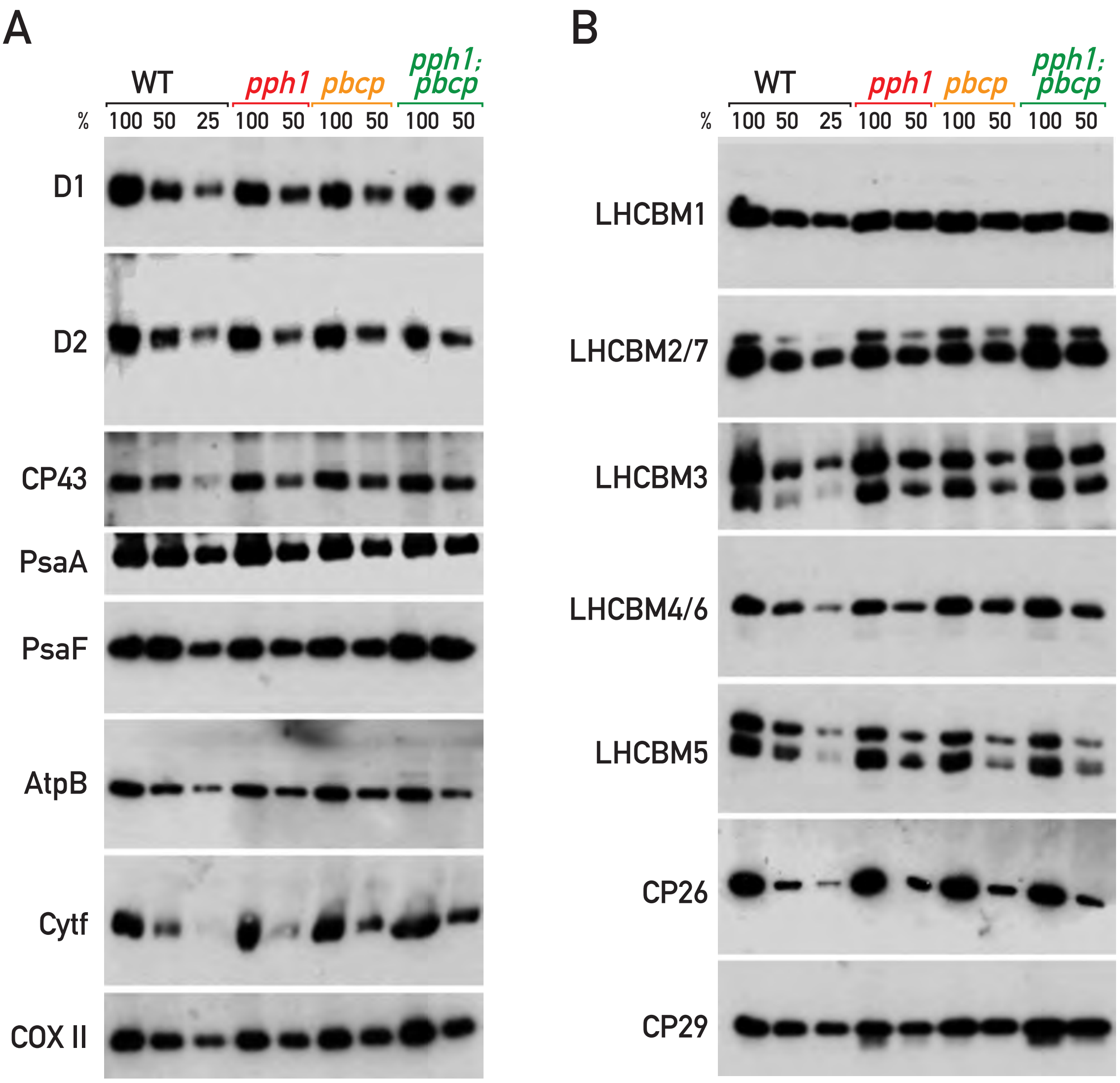


**Fig. S8. Accumulation of photosynthetic proteins in the *pph1*, *pbcp* and *pph1;pbcp* mutants**

Immunoblot with antiserum against D1, D2, CP43, PsaA, PsaF, AtpB, Cytf, COXII (mitochondrial), LHCBM1, LHCBM2/7, LHCBM3, LHCBM4/6, LHCBM5, CP26 and CP29 on total protein extracts from *pph1, pbcp, pph1;pbcp* and wild type, loaded in different amounts (100 % corresponds to 10 µg of total protein).

### Figure S9

```
type II
LHCBM5 Cre03.g156900.t1.2 ----MMLSRTVVNVQAKLTKKGGAPKKAAPASAQKKTIREKAGWWSNGGNEKLSAFYGPDRGLWLGPLS-GTTPAYLTGEFPGDYGWDSAGLSADPE

type IV
LHCBM1 Cre01.g066917.t1.1 MAFALAKSSARAAVSRRSTVKVEARRTVKP---------------ASKASTPDSFWYGPERPLFLGAFT-GEPPSYLTGEFPGDYGWDTAGLSADPE
Peptide LHCBM1                                   ARRTVKP---------------ASKASTPD

type III
LHCBM2 Cre12.g548400.t1.2 MAAIM---KSAVRSS-----VRPT-VSGRSA---------------RVVPRAAIEWYGPDRPKFLGPFSEGDTPAYLTGEFPGDYGWDTAGLSADPE
LHCBM7 Cre12.g548950.t1.2 MAAIM---KSSVRSS-----VRST-VSSRSA---------------RVVPRAAIEWYGPDRPKFLGPFSEGDTPAYLTGEFPGDYGWDTAGLSADPE
Peptide LHCBM2/7                                                                  GPDRPKFLGPFSEGD

type I
LHCBM3 Cre04.g232104.t1.1 MAFALSFSRKALQVSAKATGKKGT-GKTAAK---------------QAPASSGIEFYGPNRAKWLGPYSENATPAYLTGEFPGDYGWDTAGLSADPE
Peptide LHCBM3                                     GKTAAK---------------QAPASS

LHCBM4 Cre06.g283950.t1.2 MAFALATSRKALQVTCKATGKKTA-AK------------------AAAPKSSGVEFYGPNRAKWLGPYSENSTPAYLTGEFPGDYGWDTAGLSADPE
LHCBM6 Cre06.g285250.t1.2 MAFAL-ASRKALQVTCKATGKKTA-AK------------------AAAPKSSGVEFYGPNRAKWLGPYSENATPAYLTGEFPGDYGWDTAGLSADPE
LHCBM8 Cre06.g284250.t1.2 MAFALATSRKALQVTCKATGKKTA-AK------------------AAAPKSSGVEFYGPNRAKWLGPYSENSTPAYLTGEFPGDYGWDTAGLSADPE
Peptide LHCBM3/4/6/8                                                        AAPKSSGVEFYG

LHCBM9 Cre06.g284200.t1.2 MAFALASSRKALQVTCKATGKKTA-AK------------------AAAPKGAGIEFYGPNRAKWLGPYSENSTPAYLTGEFPGDYGWDTAGLSADPE
```

**Fig. S9. Design of peptide antigens for antisera against LHCBM isoforms.**

Alignment of LHCBM N-terminal polypeptide sequences. The predicted chloroplast transit peptides are indicated in green (Natali and Croce, 2015), and the phosphorylated Thr in red. The peptides used as the respective antigens are indicated in purple.

### Figure S10

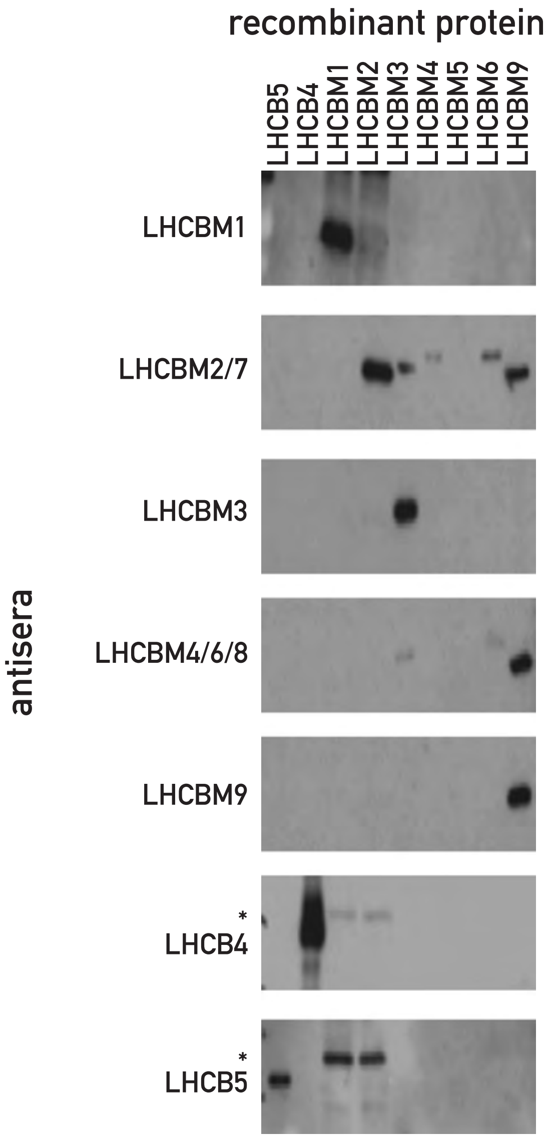


**Fig. S10. Specificity and cross-reactions of the antisera against LHCII components.**

Immunoblot with the antisera against LHCII components on total protein extracts from *E.coli* expressing the different recombinant proteins (Girolomoni *et al.* 2017). The asterisk indicates a cross-reacting band from *E.coli*, which is more prevalent in the LHCBM1 and LHCBM2/7 lanes because these recombinant proteins were expressed to lower levels so that larger amounts of the respective bacterial extracts were loaded.

**Table S1. Maximum quantum yield of PSII and chlorophyll content**

| Strain | Fv / Fm (a) | Chl tot (b) | Chl *a* / Chl *b* (b) |
|---|---|---|---|
| **wt** | 0.74 ± 0.03 | 1.86 ± 0.45 | 1.23 ± 0.22 |
| ***pph1*** | 0.74 ± 0.02 | 1.46 ± 0.24 | 1.54 ±0.48 |
| ***pbcp*** | 0.70 ± 0.04 | 1.84 ± 0.37 | 1.30 ± 0.11 |
| ***pph1;pbcp*** | 0.70 ± 0.03 | 1.66 ± 0.36 | 1.26 ±0.16 |

(a) Maximum quantum yield of PSII ($F_v/F_m = (F_m-F_o)/F_m$). Cells were grown in TAP medium under 60 µmol $m^{-2}$ $s^{-1}$ white light. Mean ± SD, n = 5

(b) Chlorophyll concentrations determined according to Porra et al. (1989). Mean ± SD, n = 5

#### Table S2. List of oligonucleotide primers

| Primer Name | Primer Sequence (5' > 3') |
|---|---|
| pPL_56(Hyg_F) | GGAACAAAAGCTGGGTACCggtacccgcttcaaatacg |
| pPL_57(Hyg_R) | CGGGCAGGTGTGTGCTCGAGctttcttgcgctatgacacttccagc |
| pFC_217 (CrPPH1_CDS_F) | cactgctactcacaacaagccgatatcATGGCACTTGCGTCGTC |
| pFC_216 (CrPPH1_CDS_R) | taggggtaggatccaagcttagatctCTTCTTGTTGAGGAAGCCGAACATGCCG |
| pFC_218 (CrPBCP_CDS_F) | cactgctactcacaacaagccgatatcATGCACACTCAACTGCTGCAAC |
| pFC_219 (CrPBCP_CDS_R) | taggggtaggatccaagcttagatctCTGGGCAGGAGCCGGG |
| pFC_85 | GGCCACAATCAGGAACTCGTC |
| pFC_92 | GCCAATGGACCAACCCAAAC |
| pFC_168 (oMJ913) | GCACCATCATGTCAAGCCT |
| pFC_169 (oMJ944) | GACGTTACAGCACACCCTTG |
| pFC_180 | TCGGTGAACCCAACCATAAT |
| pFC_181 | AAAGATGTGGGTCTGGAACG |
| pFC_109 (mid_up) | ATGGCCTGTTTCTTAGC |
| pFC_110 (mid_low) | CTACATGTGTTTCTTGAC |
| pFC_112 (fus1_up) | ATGCCTATCTTTCTCATTCT |
| pFC_113 (fus1_low) | GCAAAATACACGTCTGGAAG |
| pFC_131 | GGACCTGTACTCGGGGTCTA |
| pFC_135 | CAGTGCACGCAACGCATC |
| pFC_265 (CrPPH1_ΔcTP_CDS_F) | CATATGGCTGCAGCGGGTCCTTC |
| pFC_266 (CrPPH1_ΔcTP_CDS_R) | GTCGACTTACTTCTTGTTGAGGAAGCCGAAC |
| pFC_267 (CrPBCP_ΔcTP_CDS_F) | CATATGGGGGGAGGTTTCCTCGCC |
| pFC_268 (CrPBCP_ΔcTP_CDS_R) | GTCGACTTACTGGGCAGGAGCCGGG |
